# Distinct EEG microstate signatures across different pain types

**DOI:** 10.64898/2026.09.28.755184

**Authors:** Pouya Rabiei, Alyson Champagne, Lorie-Ève Barrette, Nadim Fakhry, Hugo Massé-Alarie, Michel-Pierre Coll

**Author notes:** **Correspondance to:** Michel-Pierre Coll, School of Psychology, Université Laval, Québec, QC, Canada.

## Abstract

Pain is associated with dynamic brain reorganization. Electroencephalographic microstates can characterize transient large-scale brain states, yet findings across pain studies are inconsistent. Two features may contribute: conditions have been studied in isolation, and analytical pipelines vary widely. We applied a harmonized analysis pipeline across three pain conditions: capsaicin-induced pain, assessed within participants against a pain-free baseline, and non-chronic and chronic back pain, each compared with matched controls. Using a five-minute resting-state electroencephalographic recordings we examined microstate topography, temporal expression and transition probabilities, and directly compared pain-related differences across pain states. We identified five microstates (A-E), corresponding to auditory, visual, salience, and attentional networks. Capsaicin pain showed increased expression of microstate C and reduced expression of microstates A, D and E. Transitions were redistributed towards microstate C, indicating a shift toward salience-related processing at the expense of sensory and attentional dynamics. In clinical back pain, effects were smaller and diverged by pain type: non-chronic back pain showed lower microstate C occurrence, opposite in direction to the capsaicin effect, whereas chronic back pain showed lower microstate D global explained variance, suggesting salience and attentional differences in clinical pain. Direct comparisons showed weaker pain-related increases in microstate C expression in both clinical back pain groups than capsaicin pain, with additional differences involving microstates A, D and E. These findings argue against a unitary EEG microstate signature of pain and instead reveal pain-type-specific patterns of large-scale brain dynamics, with microstate C providing the clearest distinction between experimental and clinical pain.

## 1. Introduction

Brain imaging studies have advanced our understanding of how pain is processed in the brain [1; 2; 6; 11; 13; 20; 37]. However, many studies have relied on static measures despite the inherently dynamic nature of brain activity and continuous reconfiguration of large-scale brain networks [6; 25]. Because pain engages distributed neural systems involved in sensation, salience, attention, and affect [3; 9; 25], characterizing these temporal dynamics may provide important insight into brain organization across pain types.

Electroencephalography (EEG) microstate analysis provides a framework for examining fast temporal brain dynamics by modeling multichannel EEG as a sequence of transient, quasi-stable topographies lasting ~60–120 ms [24; 27; 33]. A limited set of recurrent microstates, commonly labelled A-F, has been associated with large-scale networks involved in auditory and visual processing, salience/default-mode processing, and attention [17; 21; 31; 33; 38; 43]. These microstates have been associated with activity in different large-scale brain networks, including the auditory network (microstate A), the visual network (microstate B), the salience and default mode networks (microstates C and E), attentional networks (microstate D), and the anterior default mode network (microstate F) [10; 33]. Their temporal characteristics may therefore provide information about the dynamic organization of ongoing brain activity [33].

In pain, reported microstate alterations vary across pain types. Microstate C shows reduced predominance in chronic pain[17; 31] and reduced expression during sustained capsaicin pa [38], whereas high-frequency sensitization paradigms implicate microstate C alongside A, B, and E [21]. Microstate D, is likewise reduced its expression in chronic pain[17; 31] particularly at higher pain intensities, but shows longer duration during capsaicin pain [38]. These divergent patterns suggest that microstate alterations may depend on whether pain is sustained, sensitizing, or chronic.

Yet no study has directly compared microstate dynamics across pain types under matched analytical conditions, leaving unclear whether these divergent patterns reflect genuine differences between clinical and experimentally induced pain or methodological heterogeneity in preprocessing, cluster selection, template definition, and temporal metrics. Because pain types differ in duration, clinical relevance, experimental control, and underlying mechanisms, a harmonized comparison is needed to distinguish shared from type-specific microstate patterns.

Here, in three distinct pain types: non-chronic back pain (nCBP), chronic back pain (CBP), and experimentally induced capsaicin pain (Cap) each compared with an age- and sex-matched control group (−MC) or, for experimental pain, within individuals (Fig1A,) we applied a harmonized EEG microstate-analysis pipeline to examined pain-related differences in microstate topography, temporal parameters, and transition probabilities and directly compared their magnitude across pain types (Fig1B-1C). Overall, our results revealed pain-type-specific differences in microstate temporal dynamics and transitions rather than a single shared microstate signature across pain types (Fig1D).

**Fig 1.**
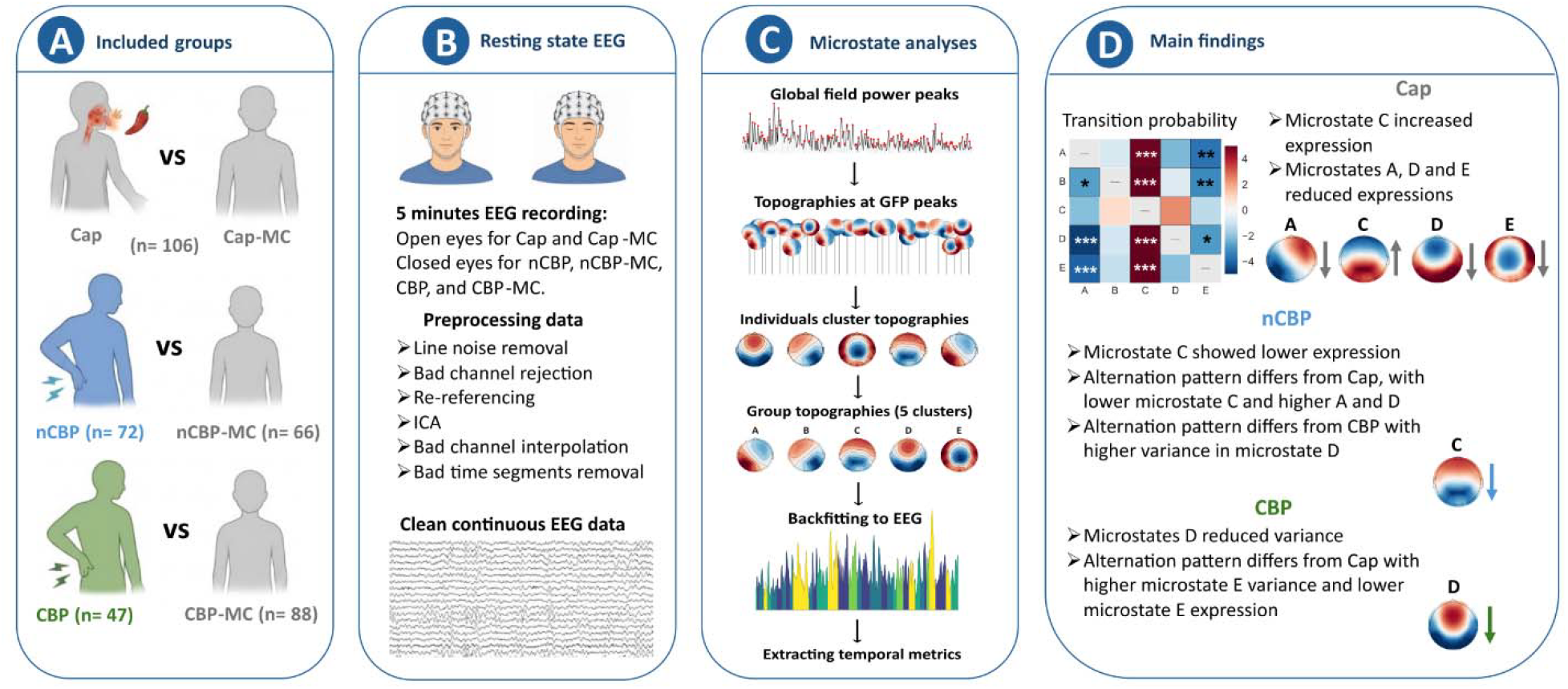
Study design, EEG microstate-analysis workflow, and principal findings across three pain types. **A:** A total of 379 participants were included across three pain–control comparisons: **B:** Five-minute EEG recordings were acquired with eyes open for the capsaicin conditions and eyes closed for the non-chronic and chronic back pain groups. EEG data underwent a harmonized preprocessing pipeline. **C:** Microstate analysis was performed by identifying global field power peaks, extracting scalp topographies at those peaks, clustering topographies at the individual and group levels, deriving five group-level microstate maps (A-E), and backfitting the resulting templates to the continuous EEG. Temporal parameters included GEV, mean duration, occurrence, and time coverage. **D:** The main findings revealed type-specific differences in microstate temporal dynamics and transitions. Arrows indicate higher or lower metrics relative to the corresponding matched control group or condition. Asterisks in the transition-probability matrix represent FDR-corrected statistical significance *** (p < 0.001), ** (p < 0.01), and * (p < 0.05). Cap, capsaicin pain; nCBP, non-chronic back pain; CBP, chronic back pain; MC, matched control/condition; EEG, electroencephalogram; GFP, global field power; GEV, global explained variance

## 2. Materials and methods

### 2.1. Participants and study design

The nCBP dataset was obtained as part of an ongoing longitudinal cohort study conducted at the Pain and Learning Lab, Université Laval, Québec, Canada. Participants with non-chronic back pain did not meet the National Institutes of Health duration and frequency criteria of chronic low back pain (pain less than three months, or if pain is more than three months, pain presents for < 50% of days in the last six months) [12]. Back pain was defined as pain in the low back area with or without leg pain above the knee limiting activities or daily routine [14]. Participants were recruited if they were aged between 18 and 60 years old and if back pain was the main complaint. Potential participants were excluded if their pain was related to specific conditions (e.g., cancer, vertebral fracture, infections, rheumatoid diseases, neuropathic pain [>4 at the DN4 questionnaire] [29]) or if they had a history of major psychiatric diseases or severe general health issues.

The nCBP-MC, Cap, and Cap-MC datasets were obtained from a separate ongoing study conducted at the Pain and Learning Lab, Université Laval, Québec, Canada, which aimed to develop EEG-based biomarkers of experimental pain perception. Capsaicin recordings in this dataset were collected during an experimental session which included other pain and aversive stimulations, including thermal pain and unpleasant auditory and taste stimulations, while resting-state recordings were collected at the beginning of the experimental session. These datasets included healthy young adults. Participants were excluded if they reported chronic or persistent pain, pregnancy or breastfeeding, a neurological or psychiatric diagnosis, use of medication affecting attention or vigilance, or the absence of normal or corrected-to-normal vision, audition, and olfaction.

CBP and CBP-MC group are openly shared (osf.io/srpbg) [16] and have been previously used in pain-related EEG studies, including a microstate study by May et al [31]. They were collected at PainLabMunich, Technical University of Munich, Munich, Germany. Inclusion criteria for the chronic back pain and control groups are described in detail in the original publications [13; 31]. Briefly, participants were recruited in the chronic back pain group if they had clinical diagnosis of chronic pain, with pain lasting at least six months, and a minimum reported average pain intensity of at least 4 of 10 during the past four weeks. Exclusion criteria included acute pain or recent changes in the pain condition during the previous three months, major neurological disorders, major psychiatric disorders other than depression, severe medical conditions, and benzodiazepine use.

### 2.2. Ethics approval and consent to participate

The nCBP dataset was approved by the Research Ethics Committee of the CIUSSS Capitale-Nationale (approval no. #2023-2682). The nCBP-MC, Cap and Cap-MC datasets were approved by Research Ethics Committee of the CIUSSS Capitale-Nationale (approval no. #2023-2487). The CBP and CBP-MC datasets were collected under ethics approval from the Medical Faculty of the Technical University of Munich, as described in the original study [13]. Written informed consent was obtained from all participants before participation. Participants included in the publicly available dataset also provided written informed consent in the original study [13].

### 2.3. Data acquisition

For the nCBP and nCBP-MC, EEG data was recorded during an eyes-closed resting-state condition while for Cap and Cap-MC, EEG was recorded during an eyes-open condition. In the Cap group, pain was induced using Tabasco Habanero sauce (7,000 Scoville units; www.tabasco.com/hot-sauces/habanero-sauce; as in Lee et al. [26]). A 0.1 mL dose of capsaicin-rich hot sauce was placed on an edible filter paper and spread over a one-cm circle. Participants then placed the side of the filter paper with the sauce on their tongue and kept their mouth closed for 30 s. Afterward, the paper was removed, and participants breathed through their mouth for one minute to allow the sauce to dry at a specific point on the tongue. They then closed their mouth for the duration of the EEG recording. In a second recording session always occurring after passively experiencing the hot sauce, the same procedure was carried out, and participants were asked to continuously rate their pain on a 0-200 visual analogue scale, with 100 being the pain threshold. Only participants that reported experiencing pain during the second recording session were included in the current analysis to ensure that the pain experience was sufficiently intense. The Cap-MC pain completed an eyes-open resting-state recording. For the nCBP, nCBP-MC, Cap, and Cap-MC datasets, EEG data were acquired using a 64-channel gel-based system (BrainVision ActiCap; Brain Products GmbH, Germany) positioned according to the extended international 10–20 system. Signals were amplified with a gain of 50,000, low pass filtered at 500 Hz and sampled at 1,000 Hz. The reference electrode was placed at FCz and the ground at FPz with impedances kept below 20 kΩ.

For the CBP and CBP-MC datasets, eyes-closed resting-state EEG recordings were selected for the present study. Full details of the recording procedures have been reported previously [31]. In brief, EEG data were recorded with 64 electrodes and a BrainAmp MR plus amplifier (Brain Products, Munich, Germany). The electrodes included all electrodes from the International 10-20 system. All EEG electrodes were referenced to electrode FCz and grounded at electrode AFz. All data were sampled at 1,000 Hz and band-pass filtered between 0.016 and 250 Hz with impedances maintained below 20 kΩ.

Across all datasets, EEG was recorded for five minutes while participants were seated comfortably and instructed to remain relaxed and awake.

### 2.4. EEG data analysis

All analyses were conducted using a standardized pipeline applied to the raw EEG recordings of each dataset.

#### 2.4.1. Preprocessing

Preprocessing of EEG data were implemented in Python using the MNE-Python package [18]. Continuous EEG data were high-pass filtered of 1 Hz for artifact identification along with a notch filter to remove low-frequency drifts and electrical line noise (60 Hz was applied for nCBP, nCBP-MC, Cap, and Cap-MC datasets and 50 Hz was applied for CBP and CBP-MC). Data were re-referenced to the common average. Noisy channels were automatically identified using the PyPREP pipeline [5], which combines multiple robust criteria, including correlation with neighboring channels, amplitude deviation, and RANSAC-based detection [22]. Identified bad channels were marked and removed from subsequent artifact-estimation steps. Independent component analysis (ICA) was performed to remove physiological and non-physiological artifacts. For ICA training, continuous raw data were segmented into non-overlapping five-s epochs, and epochs exceeding a peak-to-peak amplitude of ±300 µV were rejected. ICA was computed using the extended Infomax algorithm and trained on cleaned five-s epochs. Independent components were automatically classified using ICLabel as implemented in mne-icalabel [28]. Components assigned a probability of >0.70 to a non-brain (muscle artifact, eye blink/movement, heartbeat, channel noise, and line noise) classes were excluded. The trained ICA solution was then applied to the continuous raw data. Components classified as non-brain sources were removed, after which the identified bad channels were interpolated, and signal jumps higher than ± 100 µV were marked for rejection.

The preprocessed EEG data were screened for data quality before microstate analysis. Participants with >10 bad channels were excluded because microstate clustering relies on stable scalp topographies. This threshold corresponded to approximately 15-16% of the 64-channel montage and approximated the highest observed group-level mean plus 2 standard deviations for bad-channel count. Based on this criterion, one participant was excluded from the Cap group. Following these exclusions, the remaining cleaned continuous EEG recordings were used for microstate analysis.

#### 2.4.2. Microstate topographies

Each preprocessed EEG file was band-pass filtered between 1 and 40 Hz and downsampled to 125 Hz [10; 31]. Microstate analysis was conducted in a two-step clustering procedure using a modified k-means algorithm [36] performed separately for each group [31; 34]. Microstates analyses were performed using the pycrostates package [15] and followed general guidelines described in previous studies [24; 33] to perform microstate analysis and group comparisons of microstate characteristics. An open-source Python package was used for analyzing EEG microstates developed to work with MNE-Python [15].

First, modified k-means clustering was performed at the individual level. For each participant, EEG topographies at global field power (GFP) peaks were clustered, generating a variable number of individual-level topographies. A total of 1,000 GFP with a minimum peak distance of 10ms were randomly selected for each participant [17; 21]. GFP was defined as the spatial standard deviation of EEG voltage values across all electrodes at each time point. GFP provides a reference-independent measure of the instantaneous strength of the scalp potential field by considering all recording electrodes simultaneously [40]. EEG topographies were clustered at GFP maxima as they represent the time points of highest signal-to-noise ratio [36]. To select the optimal number of clusters, the clustering procedure was repeated for different values of k, ranging from 3 to 8. For each candidate k, the initial set of cluster topographies was randomly drawn from the selected GFP-peak maps, and the clustering was repeated 100 times with different random initializations to reduce seed-dependent variability. After fitting the model, the resulting cluster centers were treated as candidate microstate templates and spatially compared with the selected GFP-peak topographies. The templates and the observed topographies were normalized to the unit norm, and their similarity was quantified using the absolute spatial correlation, thereby ignoring polarity. Each GFP-peak topography was assigned to the cluster with the highest spatial correlation. Topographies whose maximum correlation with any template was below the threshold of 0.5 were labeled as unassigned [10; 31; 42]. This procedure was repeated across all tested values of k to evaluate the clustering solutions and identify the optimal number of microstate clusters. For each value of k, repeated clustering runs were evaluated using global explained variance (GEV) as the proportion of global variance that is explained by every microstate. The run explaining most variance was retained as the representative clustering solution for that specific k [31]. The corresponding set of cluster centers was then selected as the individual-specific microstate template set for that candidate cluster. The optimal number of clusters was identified for each individual separately based on when the increase in GEV for one number of microstates (relative to the previous number of microstates) was <5% [23].

Second, the selected individual-level microstate topographies were concatenated and submitted to group-level modified k-means clustering. Candidate solutions from 3 to 8 clusters were each repeated 200 times, retaining the highest-GEV solution for each k and applying the same < 5% relative GEV-increase criterion to select the optimal group solution. This procedure yielded a five-cluster solution for all datasets, consistent with previous pain-related microstate studies [21; 31], and the resulting maps were labeled A-E (Supplementary Fig S1 and Table S1).

#### 2.4.3. Microstates temporal metrics

The final group-level microstate templates were backfitted to the continuous EEG data of each participant separately within each group. For Cap and Cap-MC groups the topographies from both conditions were backfitted using a common pooled Cap/Cap-MC five-cluster solution, generated by entering the data into a single clustering procedure given that this was a within-subjects design [21; 33]. This ensured that farther analyses were performed with respect to the same A to E microstate labels in both paired conditions. Then, temporal segmentation was then performed using the group-specific aligned microstate map. For each participant and each microstate cluster, the resulting microstate sequence was used to compute standard temporal metrics, including mean duration, time coverage, frequency of occurrence, and GEV. The mean duration is the average time (in milliseconds) for which a microstate persists before transitioning to a different microstate. The time coverage is the percentage of total time that a microstate is present. The frequency of occurrence is the number of times that a microstate recurs per second. GEV is the percentage of global variance that is explained by every microstate.

### 2.5. Statistical analysis

All statistical analyses were performed in Python using the pingouin [41] and statsmodels [39] packages. Statistical significance was assessed using two-sided tests. For all analyses multiple comparisons were corrected using the Benjamini–Hochberg false discovery rate (FDR) procedure [4].

Group comparison analysis for A to E microstates’ maps was performed via topographic analyses of variance (TANOVAs) permutation tests, a nonparametric randomization test based on the global map dissimilarity of individual topographies [35]. Global map dissimilarity is a measure of the difference between two topographies related to the spatial correlation [35]. For each microstate cluster, global map dissimilarity was computed between the average microstate topographies of the two groups/conditions. P-value was obtained by comparing dissimilarity with a distribution of dissimilarities generated by randomly shuffling individual topographies between both groups and re-computing the dissimilarity between the center topographies of the randomized groups. The process was repeated 5,000 times. The result was a P-value per microstate, which was given by the proportion of permutations in which the dissimilarity was smaller than the dissimilarity originally observed in the data. Resulting P-values were corrected for multiple comparisons across the five microstate clusters.

Transition probabilities were computed from the backfitted microstate sequences using transition matrices with repeated consecutive states ignored. For each subject, transition probabilities were row-normalized, such that outgoing probabilities from a given microstate summed to one. Non-self transitions between microstates A to E were analyzed, yielding 20 directed transitions per comparison. For nCBP versus nCBP-MC and CBP versus CBP-MC, transition probabilities were compared using independent-samples t-tests. For Cap versus Cap-MC, transition probabilities were compared using paired-samples t-tests after matching subjects across conditions and excluding the unmatched Cap-MC subject. FDR correction was applied separately within each planned comparison across the 20 non-self transitions. Because transition probabilities are row-wise constrained, significant transition results were interpreted as evidence of redistribution of transition dynamics rather than as independent effects for each individual transition.

Temporal microstate metrics were analyzed for GEV, mean duration, occurrence, and time coverage. Because the study included both independent group comparisons (nCBP versus. nCBP-MC and CBP versus. CBP-MC) and a paired experimental pain comparison (Cap versus Cap-MC) statistical models were adapted to the structure of each comparison. For nCBP versus. nCBP-MC and CBP versus CBP-MC, omnibus mixed ANOVAs were performed separately for each temporal metric, with group as a between-subject factor and microstate cluster as a within-subject factor. For Cap versus Cap-MC, because both conditions were obtained from the same participants, omnibus repeated-measures ANOVAs were performed with condition and microstate cluster as within-subject factors.

When Group × Microstate or Condition × Microstate interactions were significant, post hoc microstate-specific tests were used to identify the microstates driving the interaction. For nCBP versus nCBP-MC and CBP versus CBP-MC, post hoc comparisons were performed using independent-samples t-tests for each metric and microstate. For Cap versus Cap-MC, post hoc comparisons were performed using paired-samples t-tests after matching subjects across conditions.

To compare whether the magnitude of pain-related differences differed between pain types, across-pair difference-of-differences analyses were performed for transition probabilities and temporal microstate metrics. For each directed non-self transition, temporal metric, and microstate, a generalized estimating equation (GEE) model with a Gaussian family and exchangeable working correlation structure was fitted with Pair, Condition, and their interaction as predictors with controlling age and sex. Pair indicated the pain type/comparison pair, and Condition indicated pain versus matched control or condition. The Pair × Condition interaction tested whether the pain-control difference differed between pain types. Pairwise difference contrasts were estimated for nCBP minus Cap, CBP minus Cap, and nCBP minus CBP, corresponding respectively to (nCBP − nCBP-MC) − (Cap − Cap-MC), (CBP − CBP-MC) − (Cap − Cap-MC), and (nCBP − nCBP-MC) − (CBP − CBP-MC). P-values were FDR-corrected across metrics, microstates, and contrasts for temporal analyses and across directed transitions and contrasts for transition analyses. GEE models were used because the combined dataset included both independent group comparisons and a paired repeated-measures comparison. Specifically, nCBP versus nCBP-MC and CBP versus CBP-MC involved independent subjects, whereas Cap versus Cap-MC involved repeated observations from the same participants. GEE provides population-average estimates while accounting for within-subject correlation through subject-level clustering [19; 30]. This approach was therefore appropriate for estimating whether the magnitude of pain-control differences differed between pain types while preserving the paired structure of the Cap comparison and maintaining unique subject identifiers for the independent nCBP and CBP comparisons.

Regression analyses were conducted to examine whether interindividual differences in pain intensity were associated with temporal microstate metrics within each pain group. Separate ordinary least-squares (OLS) models [8] were fitted for the capsaicin pain, non-chronic back pain, and chronic back pain groups for each combination of temporal metric—GEV, mean duration, occurrence, and time coverage—and microstate cluster A-E. Pain intensity was entered as the predictor and the corresponding temporal microstate metric as the outcome. Capsaicin pain ratings measured on a 0-200 scale as the current pain while clinical non-chronic and chronic back pain ratings measured on a 0-10 scale as the average of pain in the last week. Benjamini–Hochberg FDR correction was applied separately within each pain group across the 20 metric-by-microstate maps.

## 3. Results

We analyzed five-minute resting-state EEG recordings from three complementary pain types and their corresponding controls. Demographic characteristics, clinical inclusion criteria, and preprocessing-quality metrics for each group are summarized in Table 1.

**Table 1.** Demographic and preprocessing-quality data.

| Group | N | Age<br>(mean (SD)) | Sex M/F<br>(%) | Average of pain<br>intensity<br>(mean (SD)) | (Min – Max)<br>Range of pain | Bad channels<br>(mean (SD)) | Rejected ICAs<br>(mean (SD)) |
| --- | --- | --- | --- | --- | --- | --- | --- |
| Capsaicin pain |  |  |  |  |  |  |  |
| Cap | 106 | 25.21 (6.31) | 35.5/64.5 | 165.88 (30.46)<br>(0 to 200 scale) * | (102 – 200)<br>98 | 2.55 (2.39) | 14.90 (5.88) |
| Cap-MC |  |  |  | - | - | 2.56 (2.43) | 13.72 (4.75) |
| Non-chronic back pain |  |  |  |  |  |  |  |
| nCBP | 72 | 28.49 (9.97) | 48.6.4/51.4 | 4.09 (1.82)<br>(0 to 10 scale) | (2 - 8) 6 | 1.38 (1.77) | 8.72 (4.56) |
| nCBP-MC | 64 | 27.21 (6.79) | 46.9/53.1 | - | - | 1.43 (1.58) | 9.33 (4.75) |
| Chronic back pain |  |  |  |  |  |  |  |
| CBP | 47 | 58.85 (13.52) | 44.6/55.3 | 5.51 (1.61)<br>(0 to 10 scale) | (2 - 9) 7 | 2.18 (2.05) | 11.88 (4.65) |
| CBP-MC | 88 | 57.52 (14.27) | 31.8/68.2 | - | - | 1.54 (1.49) | 11.00 (5.10) |

Cap, capsaicin-induced experimental pain; nCBP, non-chronic back pain; CBP, chronic back pain; MC, matched control; SD, standard deviation; M, male; F, female ICA, independent component analysis
\* The pain scale in capsaicin-induced experimental pain was different from other groups

### 3.1. Microstate map estimation and topographic comparisons

We examined whether the spatial configuration of the identified microstates differed between pain groups and their matched controls, as marked topographic differences could influence the interpretation of subsequent temporal comparisons. Permutation-based topographic analyses of variance based on global map dissimilarity showed that topographic dissimilarity differences were group-specific: no differences survived FDR correction in the capsaicin and chronic back pain comparisons, whereas microstates B, D, and E differed between the non-chronic back pain and its matched controls (Fig2). See Supplementary Table S2 for full comparison.

**Fig 2.**
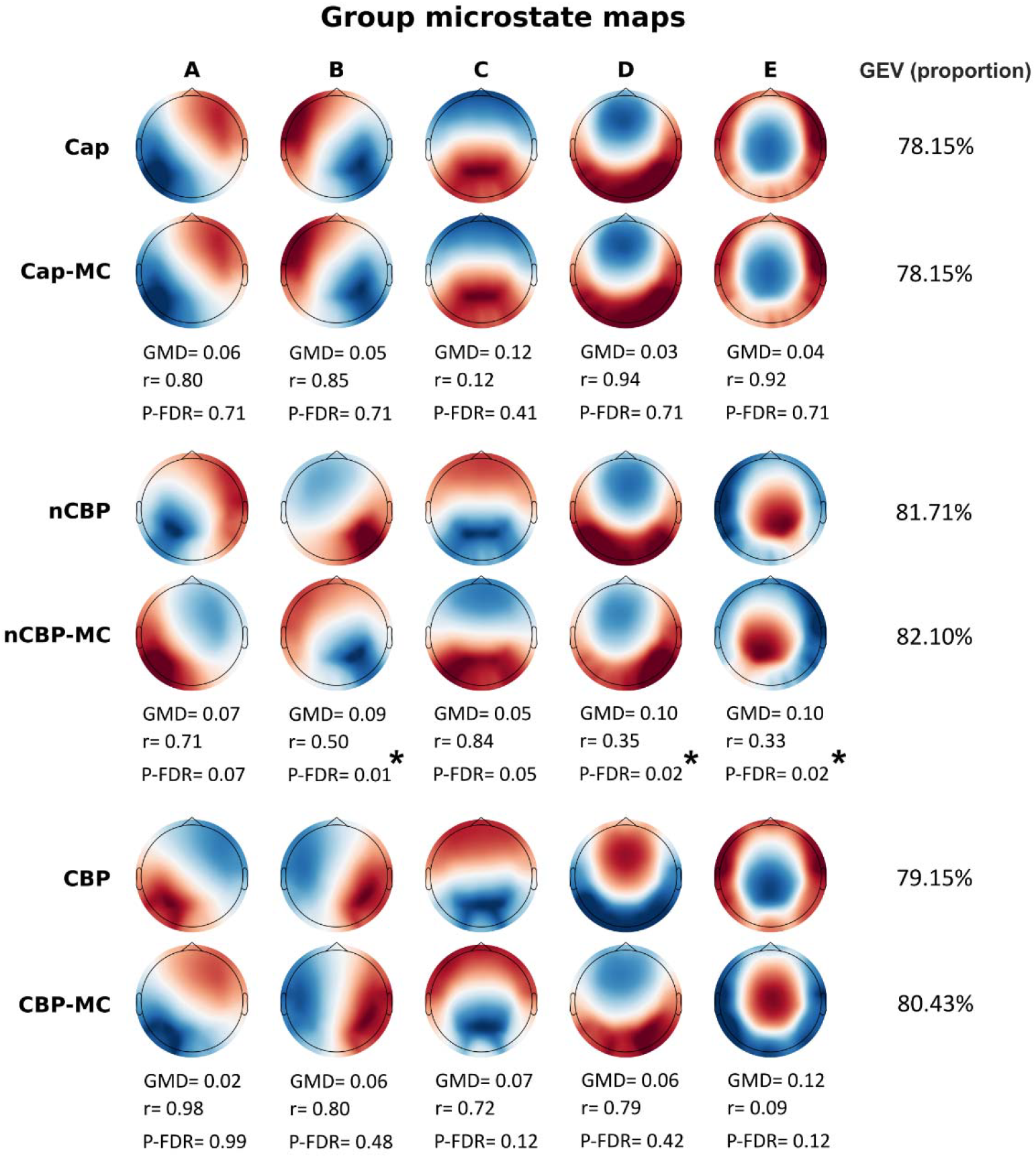
Group-level topographies of the five-microstate solutions. Microstate maps A-E are shown for each pain group and its matched control group or control condition. Microstate labels were assigned based on their correspondence with commonly reported canonical microstate clusters. GEV is reported for each group. Significant between-group topographic differences, assessed using GMD, are indicated at an FDR-corrected P value. GMD indicates the observed global map dissimilarity between group-average maps, with higher values reflecting greater topographic dissimilarity. r indicates the spatial correlation between group-average maps, with lower values indicating lower topographic similarity. P-_FDR_ values were corrected within each planned group comparison across the five microstates using the false discovery rate procedure. nCBP, non-chronic back pain; Cap, capsaicin pain; CBP, chronic back pain; MC, matched control or condition; GEV, global explained variance, GMD, global map dissimilarity. *FDR-corrected significance levels as < 0.05.

### 3.2. Transition-probability comparisons

We tested transition probabilities between microstates, excluding self-transitions, using independent samples t-tests for the non-chronic and chronic back pain comparisons and paired-samples t-tests for the within-participant capsaicin pain comparison. We additionally tested whether pain-related differences in transition probabilities differed across pain types via GEE model.

In clinical pain groups, no transitions survived FDR correction in either the non-chronic or chronic back pain comparisons. By contrast, capsaicin pain was associated with reduced transitions toward microstate A from microstates B, D, and E and toward microstate E from microstates A, B, and D. Capsaicin pain was also associated with increased transitions toward microstate C, with higher transition probabilities from microstates A, B, D, and E (Fig3A). These findings indicate that experimentally induced capsaicin pain shifted microstate sequencing toward microstate C and away from microstates A and D, whereas transition dynamics did not differ from matched condition in the two clinical back pain groups.

**Fig 3.**
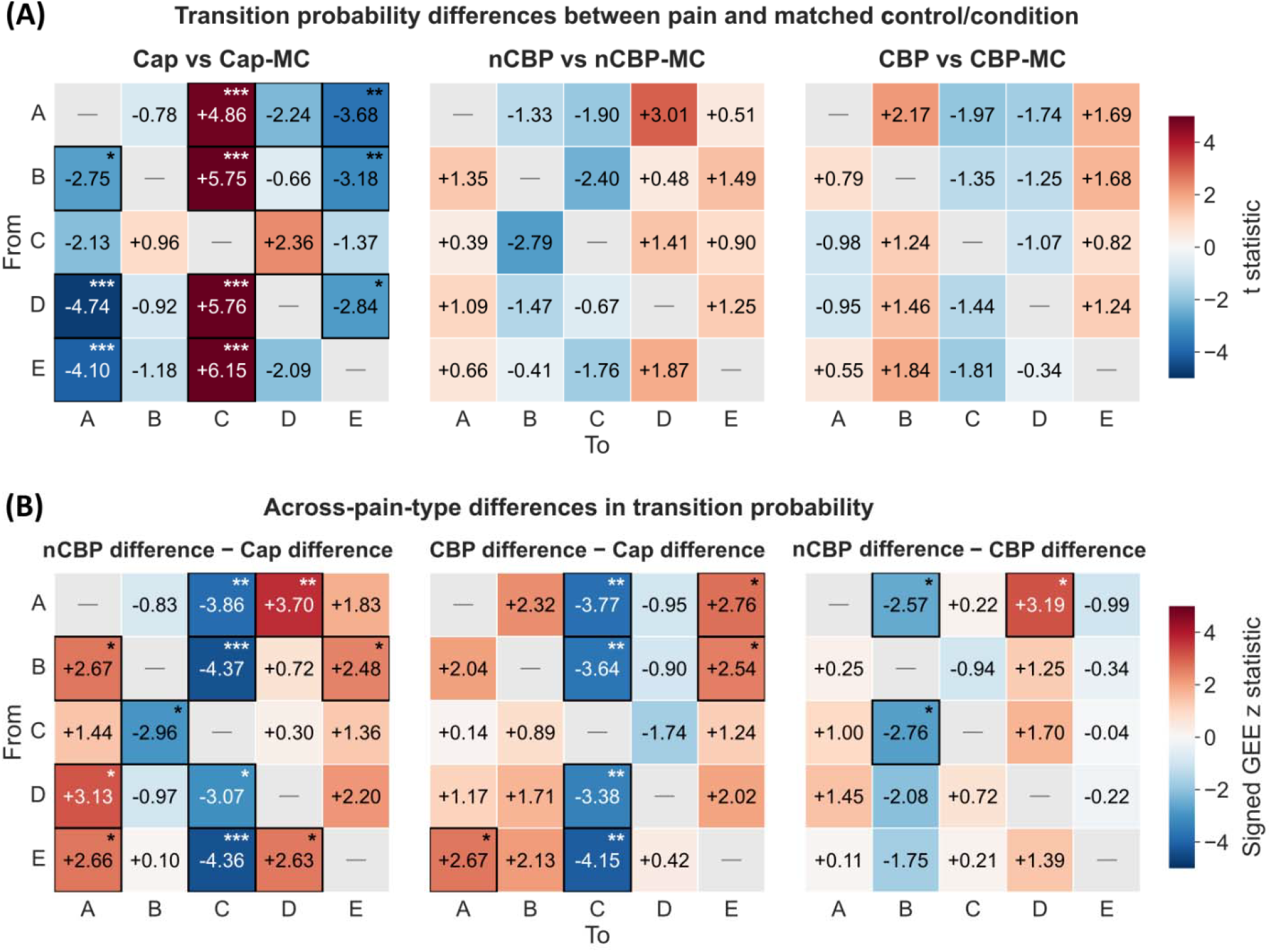
Transition-probability differences within and across pain types. **A:** t statistics for transition-probability differences between each pain group and its matched control group or control condition. Paired-samples t-tests were used for the capsaicin comparison, whereas independent-samples t-tests were used for the non-chronic and chronic back pain comparisons. **B:** Signed generalized estimating equation z statistics for differences in pain-related transition-probability effects across pain types. Positive values indicate a larger pain–control difference for the first pain type named in each contrast, whereas negative values indicate a larger difference for the second. Self-transitions were excluded. Cap, capsaicin pain; nCBP, non-chronic back pain; CBP, chronic back pain; MC, matched control; GEE, generalized estimating equation. P-value after FDR correction *** (p < 0.001), ** (p < 0.01), and * (p < 0.05)

Interaction analyses showed that pain-related transition-probability differences varied across pain types (Fig3B). The main effect observed in the capsaicin pain-control contrast was different from both the non-chronic and chronic back pain contrasts. Accordingly, both clinical groups showed more negative transition-probability differences toward microstate C from A, B, D, and E. The accompanying differences among A, B, D, and E also differed between each back-pain group and the capsaicin condition. In contrast, the non-chronic and chronic back pain effects differed for only three transitions: lower A to B and C to B as well as higher A to D transitions. Overall, the most consistent distinction between capsaicin and back pain involved lower transitions toward microstate C, whereas transition differences were more negative between the non-chronic and chronic back pain types, but not identical. Full list of statistical results for transition probability is reported in Supplementary Table S3-S

### 3.3. Microstate temporal metrics comparisons

We tested whether each pain type changed the temporal expression of individual microstates. We quantified microstate expression using four complementary metrics: GEV, mean duration, occurrence, and time coverage. We examined pain-control differences within each pain type using a 2 Group/Condition × Microstate mixed or repeated measures ANOVA model for each pain type. The full distribution of each metric in each group/condition and all statistical results are shown in Supplementary Fig S2 and Table S5.

For capsaicin pain compared with matched condition, a repeated-measures ANOVAs showed significant Condition × Microstate interactions for GEV (F(4,416)= 15.02, P-FDR< 0.001, Generalized eta-square effect size [ηg^2^] = 0.02), mean duration (F(4,416)= 20.50, P-FDR< 0.001, ηg^2^= 0.03), occurrence (F(4,416)= 13.63, P-FDR< 0.001, ηg^2^= 0.01), and time coverage (F(4,416)= 21.77, P-FDR< 0.001, ηg^2^= 0.03). Post hoc paired comparisons (Fig4A) showed that microstate C was more strongly expressed during capsaicin pain than in the no-pain condition, with higher GEV, mean duration, occurrence, and time coverage (P-FDR< 0.001). In contrast, microstates A, D and E showed reduced expression during capsaicin pain. For microstate A, the capsaicin pain condition presented with lower GEV, occurrence, and time coverage (P-FDR< 0.001). For microstate D, the capsaicin pain condition presented with lower GEV (P-FDR< 0.001), and occurrence (P-FDR= 0.03), than the no-pain condition. Finally, for microstate E, the capsaicin pain condition had lower GEV, occurrence, and time coverage (P-FDR< 0.001) than the no-pain condition.

For non-chronic back pain compared with matched controls, mixed ANOVAs showed significant Group × Microstate interaction only for occurrence (F(4,544)= 2.81, P-FDR= 0.03, Partial eta-squared effect sizes [ηp^2^]= 0.01). Post-hoc comparison showed that participants with non-chronic back pain had lower occurrence than match controls in microstate C (P-FDR< 0.001) (Fig4A).

**Fig 4.**
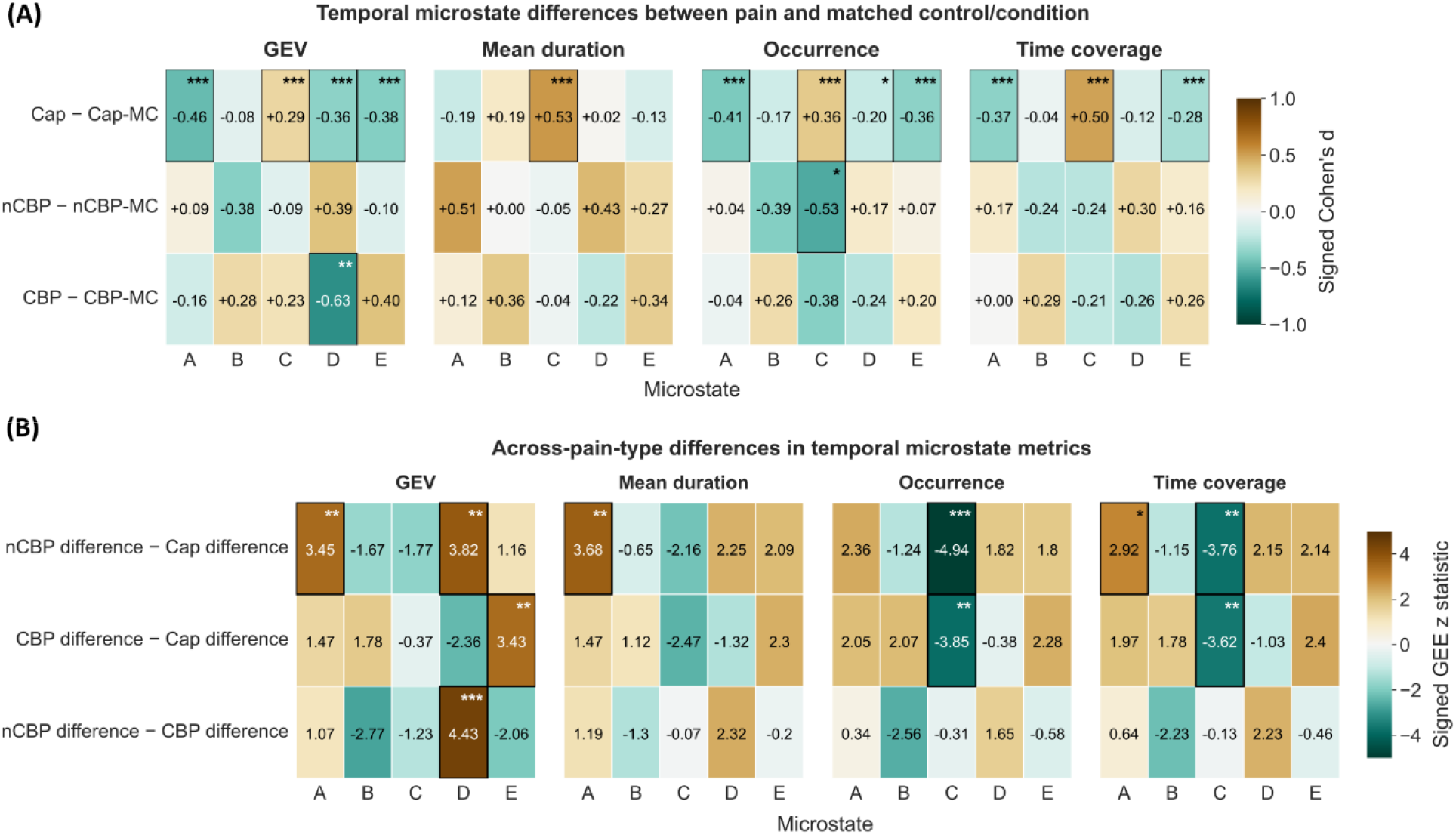
Temporal microstate differences within and across pain types. **A:** Standardized differences in global explained variance, mean duration, occurrence, and time coverage between each pain group and its matched control group or control condition. Values represent signed Cohen’s d, with positive values indicating higher metric values in the pain group and negative values indicating lower values relative to the matched control or condition. Paired-samples tests were used for the capsaicin comparison, whereas independent-samples tests were used for the non-chronic and chronic back pain comparisons. **B:** Differences in pain-control effects across pain types, estimated using generalized estimating equation models. Signed generalized estimating equation z statistics for differences in pain-related temporal metrics’ differences across pain types. Positive values indicate a larger pain-control difference for the first pain type named in each contrast, whereas negative values indicate a larger difference for the second. Cap, capsaicin pain; nCBP, non-chronic back pain; CBP, chronic back pain; MC, matched control/condition; GEE, generalized estimating equation. P-value after correction *** (< 0.001), ** (< 0.01), and * (< 0.05)

For chronic back pain compared with matched controls, mixed ANOVAs showed significant Group × Microstate interaction only for GEV (F(4,528)= 5.30, P-FDR= 0.03, ηp^2^= 0.03). Post-hoc comparison showed that participants with chronic back pain had lower GEV in microstate D (P-FDR= 0.005) compared with matched controls (Fig4A).

Then, we tested whether the magnitude of microstates differences found within each pain profile differed between pain groups. To do this, we used GEE models to estimate pairwise interaction contrasts between pain types: [nCBP − nCBP-MC] − [Cap − Cap-MC], [CBP − CBP-MC] − [Cap − Cap-MC], and [nCBP − nCBP-MC] − [CBP − CBP-MC] (Fig4B).

Compared with the capsaicin pain-control difference, the chronic back pain-control difference was negative for microstate C time coverage (difference-of-differences= −5.39, z= −3.62, P-FDR= 0.003) and occurrence (difference-of-differences = −0.28, z = −3.85, P-FDR= 0.002). In contrast, the chronic back pain difference was more positive for microstate E GEV (difference-of-differences= 0.026, z= 3.43, P-FDR= .004).

The non-chronic back pain-control difference also differed from the capsaicin contrast. For microstate C, the non-chronic back pain difference was negative for time coverage (difference-of-differences= −5.65, z= −3.76, P-FDR= 0.002) and occurrence (difference-of-differences= −0.31, z= −4.94, P-FDR< 0.001). Conversely, the non-chronic back pain difference was more positive for microstate A mean duration (difference-of-differences= 4.52, z= 3.68, P-FDR= 0.002), time coverage (difference-of-differences= 2.37, z= 2.92, P-FDR= 0.02), and GEV (difference-of-differences= 0.017, z= 3.45, P-FDR= 0.004), as well as for microstate D GEV (difference-of-differences= 0.028, z= 3.82, P-FDR= 0.002). The only significant difference between the two back pain groups was observed for microstate D GEV, for which the pain-control difference was more positive in non-chronic than chronic back pain (difference-of-differences= 0.054, z= 4.43, P-FDR < 0.001). Full comparisons are provided in supplementary Table S6. Together, these findings indicate that both back pain groups showed smaller pain-related differences in microstate C expression than the capsaicin pain. In contrast, differences involving microstates A, D, and E varied across the two clinical back pain groups, indicating more pain-type-specific differences outside microstate C.

### 3.4. Association between pain intensity with temporal microstate expression

Finally, we tested whether interindividual variation in pain intensity was associated with temporal microstate expression within each pain group. This analysis addressed a question of whether participants reporting greater pain also showed systematically higher or lower temporal metrics within a given microstate. The results showed only one negative association between pain intensity and microstate E mean duration in the Cap group survived the FDR correction: higher pain intensity was associated with shorter mean duration of microstate E in the capsaicin group (Supplementary Fig S3-S6).

## 4. Discussion

This study compared EEG microstate organization across capsaicin pain, non-chronic back pain, and chronic back pain using a harmonized analytical pipeline. Overall, the three pain types did not share a common microstate signature. Capsaicin pain showed the most extensive reorganization. Capsaicin pain produced the broadest reorganization, with greater microstate C expression, reduced expression of microstates A, D, and E, and transitions directed toward microstate C. Non-chronic back pain was characterized by lower microstate C occurrence, whereas chronic back pain showed lower microstate D global explained variance. Across-pain-type comparisons further showed that both clinical back pain groups had smaller pain-related differences in microstate C occurrence and time coverage than the capsaicin condition. These findings indicate that experimental and clinical pain involve distinct patterns of large-scale EEG dynamics rather than degrees of the same underlying microstate differences.

Capsaicin pain was associated with a coordinated increase in microstate C across temporal metrics and a redistribution of transition probabilities toward this state. This convergence suggests that microstate C became both more prominent and more central within the microstate sequence during ongoing oral capsaicin pain. Microstate C has often been associated with salience-related and default mode-oriented processes and linked to activity in the anterior insula, dorsal anterior cingulate cortex, and inferior frontal regions [7; 10; 33]. A cautious interpretation is therefore that oral capsaicin shifted large-scale brain dynamics toward the processing of an ongoing, internally generated, and behaviourally relevant nociceptive signal. However, because pain and salience were not manipulated independently, this result cannot establish that the microstate C increase was specific to nociception rather than to the broader attentional and interoceptive demands of the stimulus.

The present capsaicin findings differ from those of Qiu et al [38] who reported lower microstate C occurrence and time coverage and reduced transitions between microstate C and microstates A and B, together with greater microstate D expression, and transitions during sustained capsaicin pain. However, the authors based these results on a four-cluster solution making a direct comparison with our results difficult. To clarify this discrepancy, we ran a sensitivity analysis using a four-cluster solution comparable to that of Qiu et al., (see Supplementary Fig S7-S8). Under this solution, the microstate C effect became consistent with Qiu et al., whereas the microstate D effect remained opposite. Cluster number thus accounts for part of the discrepancy for microstate C, plausibly because topographies resolved separately as C and E in the five-cluster solution are merged differently when only four are extracted. The disagreement for microstate D, however, must reflect other differences in design, including the control condition (rest vs. placebo cream), application site, recording timing, and preprocessing. Because microstate D has been associated with attentional reorienting and externally directed control [10; 33] this divergence may be functionally meaningful: intraoral capsaicin against a resting baseline may draw attention inward toward the ongoing sensation, whereas cutaneous application against a placebo-cream comparison may sustain a more externally directed, evaluative set. Microstate C would then index the salience of the nociceptive signal itself and microstate D the attentional context surrounding it, which would make D the more protocol-dependent of the two. With only two sustained-capsaicin studies available, this remains a hypothesis for designs that manipulate attentional demands explicitly.

The non-chronic back pain findings were more selective. The only temporal difference was lower microstate C occurrence, with no transition-probability differences. This pattern contrasts with the increased microstate C expression and transitions observed during capsaicin pain. Lower occurrence indicates that microstate C appeared less frequently, but does not establish lower duration, coverage, or overall contribution. One possible interpretation is that recent-onset clinical pain does not produce the same repeated recruitment of salience-and default mode-related configurations as an immediate experimental stimulus. Capsaicin pain was novel, intense, and continuously present during recording, whereas non-chronic back pain was part of an ongoing clinical context. The non-chronic comparison also showed topographic differences in microstates B, D, and E. These spatial differences limit purely temporal interpretations because identically labelled maps may not represent fully equivalent configurations across groups. The findings therefore support a distinct non-chronic back pain profile, but not the view that there is a continuum from experimental to chronic back pain, while non-chronic back pain would represent an intermediate stage of this continuum.

In chronic back pain, microstate D showed significantly lower global explained variance, while the other microstate D metrics were numerically lower but did not survive correction for multiple comparisons his constitutes a partial replication of May et al [31] who analysed the same recordings and reported a less prominent microstate D across temporal measures using Bayesian inference. The direction of effect agreed across all microstate D metrics, but only GEV showed both a significant difference and strong evidence against the null in the present analysis; Bayes factors for the remaining metrics were largely anecdotal. Because the participants and recordings are identical and both analyses included Bayesian inference, the weaker evidential support obtained here cannot be attributed to sampling, acquisition, or inferential framework, and is therefore attributable to the analysis pipeline itself. This directly illustrates the analytical sensitivity that motivated the present harmonized approach and indicates that the consistency of a microstate finding across studies cannot be assessed independently of the pipeline that produced it. Cumulative progress will therefore depend on convergence toward a standardized analytical framework, and, until such standards exist, on sensitivity analyses across plausible pipeline choices alongside sharing of topographies and code.

Microstate D has been associated with the dorsal attention network, frontoparietal activity, attentional switching, and reorientation [10; 33]. May et al. proposed that reduced microstate D expression may relate to attentional difficulties reported in chronic pain, where ongoing pain competes with other information for limited cognitive resources [31]. The present reduction in microstate D GEV is compatible with this account and may indicate a weaker contribution of an attention-related configuration to resting-state activity. However, attentional performance was not measured, and lower GEV does not by itself demonstrate impaired cognitive control.

The direct comparisons clarify that the capsaicin pattern was not simply a stronger version of the clinical pain patterns. Both back pain groups showed smaller pain-related differences in microstate C occurrence and time coverage than the capsaicin condition. They also differed from capsaicin pain in selected metrics involving microstates A, D, and E. In contrast, the two back pain groups differed from one another in relatively few measures, most clearly in microstate D GEV. Transition analyses showed a similar organization: both clinical groups differed from capsaicin pain for transitions toward microstate C, whereas their transition profiles were more similar to each other. These results identify microstate C dynamics as the clearest distinction between experimental capsaicin pain and clinical back pain, while microstates A, D, and E captured more condition-specific differences.

More broadly, these findings argue against treating any individual microstate cluster as a general EEG marker of pain. Microstate differences appear to depend on pain type, experimental context, recording condition, and analytical choices. This conclusion is consistent with the broader methodological concern that functional interpretations of canonical microstates should rely on convergent evidence rather than fixed labels alone [32]. The limited association with pain intensity further suggests that the observed group differences were not simple linear markers of subjective pain severity. However, single pain ratings may not adequately represent the fluctuating and multidimensional nature of pain.

A major strength of this study was the application of the same preprocessing, clustering, backfitting, and statistical framework across three complementary pain conditions. Matched controls reduced the influence of dataset-specific acquisition characteristics on within-pair comparisons. Nevertheless, the datasets differed in recruitment, recording condition, pain model, and geographic setting. Capsaicin recordings were eyes open, whereas clinical recordings were eyes closed. The capsaicin contrast was within-participant, whereas clinical contrasts were between-participant with unequal and, for chronic back pain, comparatively small groups. Differences in pain-related effects across types may therefore reflect statistical sensitivity as well as underlying phenomena, and the absence of clinical-group effects should not be interpreted as evidence of equivalence. Finally, group-specific template estimation and topographic differences in the nCBP comparison limit strict equivalence of identically labelled microstates across datasets. The mechanistic and clinical significance of these pain-type-specific patterns remains to be established. Longitudinal and intervention studies should determine whether microstate dynamics reflect underlying pain-processing mechanisms, have prognostic value for pain persistence or recovery, or predict treatment response.

In conclusion, pain was associated with type-specific rather than uniform EEG microstate reorganization. Capsaicin pain showed a broad shift toward microstate C, non-chronic back pain showed reduced microstate C occurrence, and chronic back pain showed reduced microstate D global explained variance. The chronic finding partially replicated prior work, whereas the capsaicin pattern differed substantially from earlier evidence. Together, the results show that experimental and clinical pain engage distinct forms of large-scale brain dynamics and caution against proposing a universal microstate signature of pain. Establishing whether pain-related microstate differences generalize will require standardized and transparently reported pipelines, including cluster-number selection, template definition, backfitting procedure, and the metrics reported, so that findings across pain models can be compared on common terms.

## Data availability

The data related to matched control of the non-chronic back pain and capsaicin pain will be publicly available after acceptance. Clinical non-chronic back pain data cannot be openly shared but will be available upon request.

## Code availability

Detailed code is available at GitHub:

https://github.com/pouya-rabiei/EEG_microstate

## Author contributions

P.R. and M.P.C. created the idea of the project, designed the study, and interpreted the results. P.R. conducted the statistical analyses and drafted statistical analysis aspects of the project. M.P.C. supervised the analytical design. M.P.C. and H.M.A. supervised the project. A.C., L.E.B., and N.F. contributed to data collection. PR drafted the manuscript with revision and input from M.P.C. and H.M.A. All authors reviewed and approved the final manuscript.

## Acknowledgement

We thank all participants for their contributions. We also thank Cristina Gil Ávila, Elisabeth S. May, and Markus Ploner for making the chronic pain EEG dataset publicly available.

