## Supplementary materials for "Distinct EEG microstate signatures across different pain types"

**Methods**

**Table S1.** TANOVA-based comparisons of microstate topographies across all datasets

| **Comparison** | **Microstate** | **GMD** | **r** | **P-FDR** |
| --- | --- | --- | --- | --- |
| Cap vs Cap-MC | A | 0.06 | 0.80 | 0.71 |
| Cap vs Cap-MC | B | 0.05 | 0.85 | 0.71 |
| Cap vs Cap-MC | C | 0.12 | 0.12 | 0.41 |
| Cap vs Cap-MC | D | 0.03 | 0.94 | 0.71 |
| Cap vs Cap-MC | E | 0.04 | 0.92 | 0.71 |
| nCBP vs nCBP-MC | A | 0.07 | 0.71 | 0.07 |
| nCBP vs nCBP-MC | B | 0.09 | 0.50 | **0.01** |
| nCBP vs nCBP-MC | C | 0.05 | 0.84 | 0.05 |
| nCBP vs nCBP-MC | D | 0.10 | 0.35 | **0.02** |
| nCBP vs nCBP-MC | E | 0.10 | 0.33 | **0.02** |
| CBP vs CBP-MC | A | 0.02 | 0.98 | 0.99 |
| CBP vs CBP-MC | B | 0.06 | 0.80 | 0.48 |
| CBP vs CBP-MC | C | 0.07 | 0.72 | 0.12 |
| CBP vs CBP-MC | D | 0.06 | 0.79 | 0.42 |
| CBP vs CBP-MC | E | 0.12 | 0.09 | 0.12 |

GMD indicates the observed global map dissimilarity between group-average maps, with higher values reflecting greater topographic dissimilarity. r indicates the spatial correlation between group-average maps, with lower values indicating lower topographic similarity. P-_FDR_ values were corrected within each planned group comparison across the five microstates using the false discovery rate procedure. Significant FDR-corrected results indicate microstates whose scalp topographies differed between groups. Marked bold indicating FDR-corrected significance levels as < 0.05. Cap, capsaicin pain; nCBP, non-chronic back pain; CBP, chronic back pain; MC, matched control

**Table S2.** Comparisons of microstate transition probabilities across pain and control

| **Comparison**  **(group 1 vs group 2)** | **Transition** | **Mean group 1** | **Mean group 2** | **t value** | **cohen_d** | **CI95%** | **P-FDR** |
| --- | --- | --- | --- | --- | --- | --- | --- |
| Cap vs Cap-MC | A to C | 0.29 | 0.26 | 4.86 | 0.41 | [0.02, 0.04] | **< 0.001** |
| Cap vs Cap-MC | A to E | 0.23 | 0.24 | -3.68 | 0.33 | [-0.02, -0.01] | **0.001** |
| Cap vs Cap-MC | B to A | 0.21 | 0.23 | -2.75 | 0.24 | [-0.03, 0.00] | **0.01** |
| Cap vs Cap-MC | B to C | 0.29 | 0.26 | 5.75 | 0.46 | [0.02, 0.04] | **< 0.001** |
| Cap vs Cap-MC | B to E | 0.23 | 0.24 | -3.18 | 0.26 | [-0.02, 0.00] | **0.004** |
| Cap vs Cap-MC | D to A | 0.21 | 0.23 | -4.74 | 0.41 | [-0.03, -0.01] | **< 0.001** |
| Cap vs Cap-MC | D to C | 0.33 | 0.29 | 5.76 | 0.47 | [0.02, 0.05] | **< 0.001** |
| Cap vs Cap-MC | D to E | 0.25 | 0.26 | -2.84 | 0.22 | [-0.02, 0.00] | **0.01** |
| Cap vs Cap-MC | E to A | 0.2 | 0.22 | -4.1 | 0.38 | [-0.03, -0.01] | **< 0.001** |
| Cap vs Cap-MC | E to C | 0.31 | 0.28 | 6.15 | 0.46 | [0.02, 0.05] | **< 0.001** |
| Cap vs Cap-MC | A to D | 0.27 | 0.28 | -2.24 | 0.21 | [-0.02, 0.00] | 0.056 |
| Cap vs Cap-MC | C to D | 0.3 | 0.29 | 2.36 | 0.23 | [0.00, 0.02] | 0.057 |
| Cap vs Cap-MC | A to B | 0.22 | 0.22 | -0.78 | 0.06 | [-0.01, 0.01] | 0.46 |
| Cap vs Cap-MC | B to D | 0.27 | 0.27 | -0.66 | 0.06 | [-0.01, 0.01] | 0.51 |
| Cap vs Cap-MC | C to A | 0.21 | 0.22 | -2.13 | 0.2 | [-0.02, 0.00] | 0.06 |
| Cap vs Cap-MC | C to B | 0.22 | 0.21 | 0.96 | 0.09 | [0.00, 0.01] | 0.40 |
| Cap vs Cap-MC | C to E | 0.27 | 0.27 | -1.37 | 0.11 | [-0.02, 0.00] | 0.23 |
| Cap vs Cap-MC | D to B | 0.21 | 0.22 | -0.92 | 0.08 | [-0.01, 0.00] | 0.40 |
| Cap vs Cap-MC | E to B | 0.2 | 0.21 | -1.18 | 0.11 | [-0.01, 0.00] | 0.30 |
| Cap vs Cap-MC | E to D | 0.28 | 0.29 | -2.09 | 0.2 | [-0.02, 0.00] | 0.06 |
| nCBP vs nCBP-MC | A to B | 0.24 | 0.25 | -1.33 | 0.22 | [-0.03, 0.01] | 0.34 |
| nCBP vs nCBP-MC | A to C | 0.25 | 0.27 | -1.9 | 0.32 | [-0.05, 0.00] | 0.26 |
| nCBP vs nCBP-MC | A to D | 0.28 | 0.25 | 3.01 | 0.51 | [0.01, 0.05] | 0.06 |
| nCBP vs nCBP-MC | A to E | 0.24 | 0.23 | 0.51 | 0.09 | [-0.02, 0.03] | 0.69 |
| nCBP vs nCBP-MC | B to A | 0.22 | 0.21 | 1.35 | 0.23 | [0.00, 0.03] | 0.34 |
| nCBP vs nCBP-MC | B to C | 0.25 | 0.29 | -2.4 | 0.41 | [-0.06, -0.01] | 0.12 |
| nCBP vs nCBP-MC | B to D | 0.28 | 0.28 | 0.48 | 0.08 | [-0.02, 0.02] | 0.69 |
| nCBP vs nCBP-MC | B to E | 0.24 | 0.23 | 1.49 | 0.25 | [-0.01, 0.04] | 0.34 |
| nCBP vs nCBP-MC | C to A | 0.21 | 0.21 | 0.39 | 0.07 | [-0.01, 0.02] | 0.69 |
| nCBP vs nCBP-MC | C to B | 0.24 | 0.27 | -2.79 | 0.48 | [-0.05, -0.01] | 0.06 |
| nCBP vs nCBP-MC | C to D | 0.3 | 0.28 | 1.41 | 0.24 | [-0.01, 0.04] | 0.34 |
| nCBP vs nCBP-MC | C to E | 0.25 | 0.24 | 0.9 | 0.15 | [-0.01, 0.03] | 0.53 |
| nCBP vs nCBP-MC | D to A | 0.23 | 0.22 | 1.09 | 0.19 | [-0.01, 0.03] | 0.43 |
| nCBP vs nCBP-MC | D to B | 0.25 | 0.26 | -1.47 | 0.25 | [-0.03, 0.01] | 0.34 |
| nCBP vs nCBP-MC | D to C | 0.28 | 0.29 | -0.67 | 0.11 | [-0.04, 0.02] | 0.64 |
| nCBP vs nCBP-MC | D to E | 0.25 | 0.23 | 1.25 | 0.21 | [-0.01, 0.04] | 0.36 |
| nCBP vs nCBP-MC | E to A | 0.22 | 0.22 | 0.66 | 0.11 | [-0.01, 0.02] | 0.64 |
| nCBP vs nCBP-MC | E to B | 0.24 | 0.24 | -0.41 | 0.07 | [-0.02, 0.01] | 0.69 |
| nCBP vs nCBP-MC | E to C | 0.25 | 0.27 | -1.76 | 0.3 | [-0.04, 0.00] | 0.27 |
| nCBP vs nCBP-MC | E to D | 0.28 | 0.27 | 1.87 | 0.32 | [0.00, 0.04] | 0.26 |
| CBP vs CBP-MC | A to B | 0.25 | 0.22 | 2.17 | 0.38 | [0.00, 0.05] | 0.27 |
| CBP vs CBP-MC | A to C | 0.26 | 0.28 | -1.97 | 0.34 | [-0.05, 0.00] | 0.27 |
| CBP vs CBP-MC | A to D | 0.25 | 0.27 | -1.74 | 0.3 | [-0.05, 0.00] | 0.27 |
| CBP vs CBP-MC | A to E | 0.24 | 0.22 | 1.69 | 0.31 | [0.00, 0.05] | 0.27 |
| CBP vs CBP-MC | B to A | 0.24 | 0.23 | 0.79 | 0.14 | [-0.01, 0.03] | 0.48 |
| CBP vs CBP-MC | B to C | 0.25 | 0.27 | -1.35 | 0.25 | [-0.04, 0.01] | 0.34 |
| CBP vs CBP-MC | B to D | 0.25 | 0.27 | -1.25 | 0.21 | [-0.04, 0.01] | 0.34 |
| CBP vs CBP-MC | B to E | 0.26 | 0.23 | 1.68 | 0.32 | [0.00, 0.05] | 0.27 |
| CBP vs CBP-MC | C to A | 0.22 | 0.23 | -0.98 | 0.17 | [-0.03, 0.01] | 0.43 |
| CBP vs CBP-MC | C to B | 0.24 | 0.22 | 1.24 | 0.22 | [-0.01, 0.04] | 0.34 |
| CBP vs CBP-MC | C to D | 0.29 | 0.3 | -1.07 | 0.19 | [-0.05, 0.02] | 0.41 |
| CBP vs CBP-MC | C to E | 0.26 | 0.25 | 0.82 | 0.14 | [-0.02, 0.04] | 0.48 |
| CBP vs CBP-MC | D to A | 0.22 | 0.23 | -0.95 | 0.15 | [-0.03, 0.01] | 0.43 |
| CBP vs CBP-MC | D to B | 0.24 | 0.22 | 1.46 | 0.25 | [-0.01, 0.04] | 0.34 |
| CBP vs CBP-MC | D to C | 0.29 | 0.31 | -1.44 | 0.24 | [-0.06, 0.01] | 0.34 |
| CBP vs CBP-MC | D to E | 0.26 | 0.24 | 1.24 | 0.22 | [-0.01, 0.05] | 0.34 |
| CBP vs CBP-MC | E to A | 0.22 | 0.21 | 0.55 | 0.1 | [-0.01, 0.02] | 0.61 |
| CBP vs CBP-MC | E to B | 0.25 | 0.23 | 1.84 | 0.33 | [0.00, 0.05] | 0.27 |
| CBP vs CBP-MC | E to C | 0.27 | 0.29 | -1.81 | 0.32 | [-0.05, 0.00] | 0.27 |
| CBP vs CBP-MC | E to D | 0.27 | 0.27 | -0.34 | 0.06 | [-0.03, 0.02] | 0.74 |

Cap, capsaicin pain; nCBP, non-chronic back pain; Cap, capsaicin pain; CBP, chronic back pain; MC, matched control. Marked bold indicating FDR-corrected significance levels as < 0.05

**Table S3.** Comparison differences of microstate transition probabilities alternation across pain groups

| **Contrast** | **Transition** | **Estimate difference of differences** | **z value** | **CI95%** | **P-FDR** |
| --- | --- | --- | --- | --- | --- |
| nCBP minus Cap difference | A to B | -0.01 | -0.83 | [-0.03, 0.01] | 0.52 |
| nCBP minus Cap difference | A to C | -0.05 | -3.86 | [-0.08, -0.02] | **< 0.001** |
| nCBP minus Cap difference | A to D | 0.04 | 3.7 | [0.02, 0.06] | **< 0.001** |
| nCBP minus Cap difference | A to E | 0.02 | 1.83 | [0.00, 0.04] | 0.14 |
| nCBP minus Cap difference | B to A | 0.03 | 2.67 | [0.01, 0.04] | **0.03** |
| nCBP minus Cap difference | B to C | -0.06 | -4.37 | [-0.09, -0.03] | **< 0.001** |
| nCBP minus Cap difference | B to D | 0.01 | 0.72 | [-0.01, 0.03] | 0.58 |
| nCBP minus Cap difference | B to E | 0.03 | 2.48 | [0.01, 0.05] | **0.04** |
| nCBP minus Cap difference | C to A | 0.01 | 1.44 | [0.00, 0.03] | 0.26 |
| nCBP minus Cap difference | C to B | -0.03 | -2.96 | [-0.06, -0.01] | **0.02** |
| nCBP minus Cap difference | C to D | 0 | 0.3 | [-0.02, 0.03] | 0.88 |
| nCBP minus Cap difference | C to E | 0.02 | 1.36 | [-0.01, 0.04] | 0.29 |
| nCBP minus Cap difference | D to A | 0.03 | 3.13 | [0.01, 0.05] | **0.01** |
| nCBP minus Cap difference | D to B | -0.01 | -0.97 | [-0.03, 0.01] | 0.48 |
| nCBP minus Cap difference | D to C | -0.05 | -3.07 | [-0.08, -0.02] | **0.01** |
| nCBP minus Cap difference | D to E | 0.03 | 2.2 | [0.00, 0.05] | 0.07 |
| nCBP minus Cap difference | E to A | 0.03 | 2.66 | [0.01, 0.04] | **0.03** |
| nCBP minus Cap difference | E to B | 0 | 0.1 | [-0.02, 0.02] | 0.94 |
| nCBP minus Cap difference | E to C | -0.05 | -4.36 | [-0.08, -0.03] | **0.00** |
| nCBP minus Cap difference | E to D | 0.03 | 2.63 | [0.01, 0.05] | **0.03** |
| CBP minus Cap difference | A to C | -0.05 | -3.77 | [-0.08, -0.03] | **< 0.001** |
| CBP minus Cap difference | A to E | 0.04 | 2.76 | [0.01, 0.06] | **0.03** |
| CBP minus Cap difference | B to C | -0.05 | -3.64 | [-0.07, -0.02] | **< 0.001** |
| CBP minus Cap difference | B to E | 0.03 | 2.54 | [0.01, 0.06] | **0.03** |
| CBP minus Cap difference | D to C | -0.06 | -3.38 | [-0.10, -0.03] | **0.01** |
| CBP minus Cap difference | E to A | 0.02 | 2.67 | [0.01, 0.04] | **0.03** |
| CBP minus Cap difference | E to C | -0.06 | -4.15 | [-0.09, -0.03] | **< 0.001** |
| CBP minus Cap difference | A to B | 0.03 | 2.32 | [0.00, 0.06] | 0.06 |
| CBP minus Cap difference | A to D | -0.01 | -0.95 | [-0.04, 0.01] | 0.48 |
| CBP minus Cap difference | B to A | 0.02 | 2.04 | [0.00, 0.04] | 0.10 |
| CBP minus Cap difference | B to D | -0.01 | -0.9 | [-0.03, 0.01] | 0.49 |
| CBP minus Cap difference | C to A | 0 | 0.14 | [-0.02, 0.02] | 0.94 |
| CBP minus Cap difference | C to B | 0.01 | 0.89 | [-0.01, 0.04] | 0.49 |
| CBP minus Cap difference | C to D | -0.03 | -1.74 | [-0.07, 0.00] | 0.16 |
| CBP minus Cap difference | C to E | 0.02 | 1.24 | [-0.01, 0.05] | 0.34 |
| CBP minus Cap difference | D to A | 0.01 | 1.17 | [-0.01, 0.03] | 0.38 |
| CBP minus Cap difference | D to B | 0.02 | 1.71 | [0.00, 0.05] | 0.17 |
| CBP minus Cap difference | D to E | 0.03 | 2.02 | [0.00, 0.06] | 0.10 |
| CBP minus Cap difference | E to B | 0.03 | 2.13 | [0.00, 0.05] | 0.08 |
| CBP minus Cap difference | E to D | 0.01 | 0.42 | [-0.02, 0.03] | 0.81 |
| nCBP minus CBP difference | A to B | -0.04 | -2.57 | [-0.07, -0.01] | **0.03** |
| nCBP minus CBP difference | A to C | 0 | 0.22 | [-0.03, 0.04] | 0.89 |
| nCBP minus CBP difference | A to D | 0.05 | 3.19 | [0.02, 0.08] | **0.01** |
| nCBP minus CBP difference | A to E | -0.02 | -0.99 | [-0.05, 0.02] | 0.47 |
| nCBP minus CBP difference | B to A | 0 | 0.25 | [-0.02, 0.03] | 0.89 |
| nCBP minus CBP difference | B to C | -0.02 | -0.94 | [-0.05, 0.02] | 0.48 |
| nCBP minus CBP difference | B to D | 0.02 | 1.25 | [-0.01, 0.05] | 0.34 |
| nCBP minus CBP difference | B to E | -0.01 | -0.34 | [-0.04, 0.03] | 0.87 |
| nCBP minus CBP difference | C to A | 0.01 | 1 | [-0.01, 0.04] | 0.47 |
| nCBP minus CBP difference | C to B | -0.05 | -2.76 | [-0.08, -0.01] | **0.03** |
| nCBP minus CBP difference | C to D | 0.04 | 1.7 | [-0.01, 0.08] | 0.17 |
| nCBP minus CBP difference | C to E | 0 | -0.04 | [-0.04, 0.03] | 0.97 |
| nCBP minus CBP difference | D to A | 0.02 | 1.45 | [-0.01, 0.04] | 0.26 |
| nCBP minus CBP difference | D to B | -0.03 | -2.08 | [-0.06, 0.00] | 0.09 |
| nCBP minus CBP difference | D to C | 0.02 | 0.72 | [-0.03, 0.06] | 0.58 |
| nCBP minus CBP difference | D to E | 0 | -0.22 | [-0.04, 0.03] | 0.89 |
| nCBP minus CBP difference | E to A | 0 | 0.11 | [-0.02, 0.02] | 0.94 |
| nCBP minus CBP difference | E to B | -0.03 | -1.75 | [-0.06, 0.00] | 0.16 |
| nCBP minus CBP difference | E to C | 0 | 0.21 | [-0.03, 0.04] | 0.89 |
| nCBP minus CBP difference | E to D | 0.02 | 1.39 | [-0.01, 0.05] | 0.28 |

Cap, capsaicin pain; nCBP, non-chronic back pain; CBP, chronic back pain; MC, matched control. Marked bold indicating FDR-corrected significance levels as < 0.05

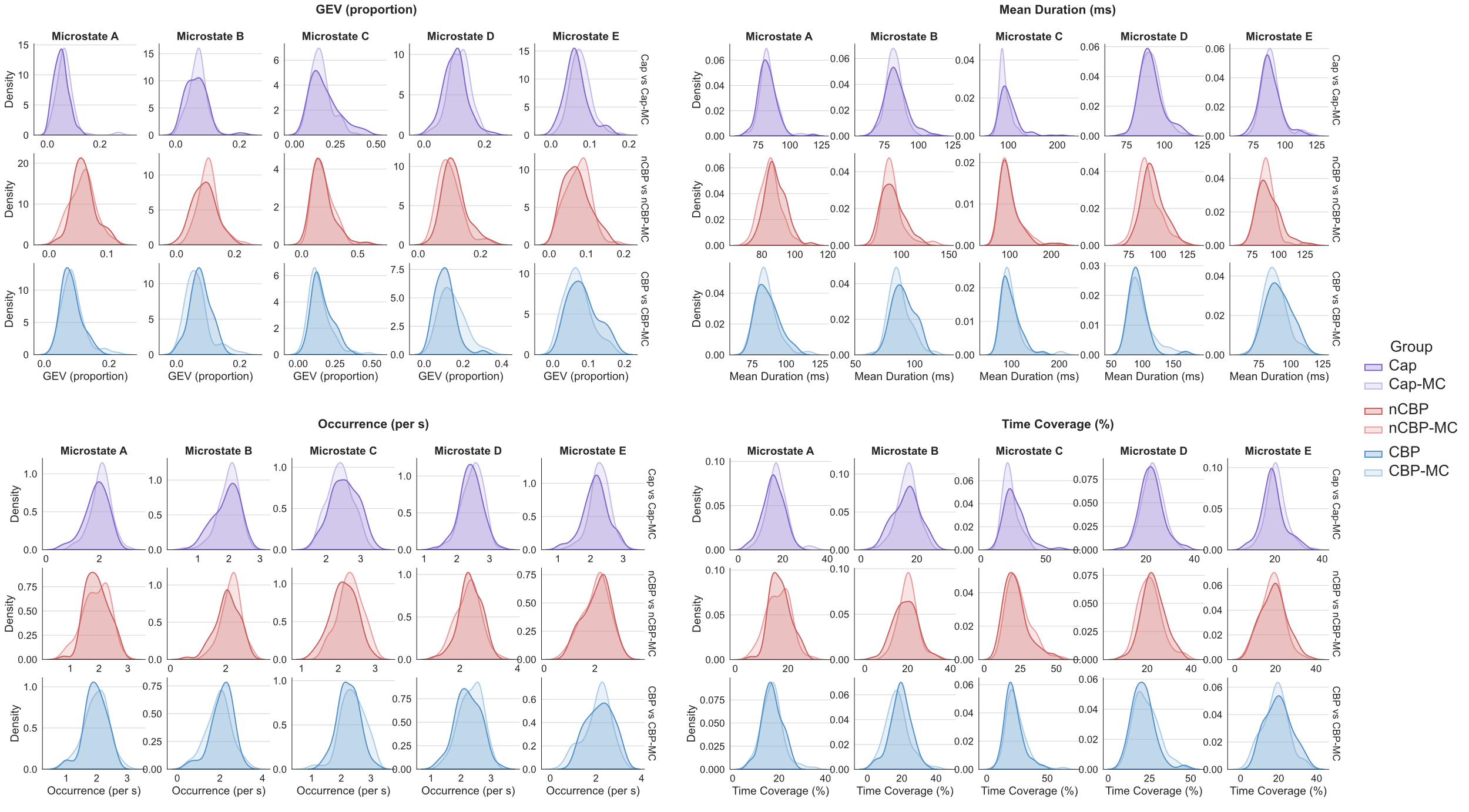

**Figure S1. Distribution of temporal microstate metrics across, capsaicin pain, non-chronic back pain, chronic back pain, and their matched control conditions.** Kernel density plots show the distributions of global explained variance, mean duration, occurrence, and time coverage for microstates A–E. Rows represent the three planned comparisons: capsaicin pain versus the matched no-pain condition, non-chronic back pain versus matched controls, and chronic back pain versus matched controls. Within each panel, darker curves indicate the pain group or condition, and lighter curves indicate the corresponding matched control group or condition. Capsaicin pain produced the broadest reorganization of temporal microstate dynamics. Relative to the no-pain condition, capsaicin pain was associated with greater expression of microstate C, reflected by higher global explained variance, mean duration, occurrence, and time coverage. In contrast, microstates D and E showed lower expression. Non-chronic back pain showed lower microstate C occurrence relative to matched controls. Chronic back pain showed lower global explained variance redistribution in microstate D. No significant interactions were observed for the other temporal metrics. Cap, capsaicin pain; nCBP, non-chronic back pain; CBP, chronic back pain; MC, matched control.

**Table S4.** Comparisons of differences of temporal microstate metrics across all datasets

| **Comparison**  **(group 1 vs group 2)** | **Microstate** | **Temporal metric** | **Mean group 1** | **Mean group 2** | **t value** | **cohen_d** | **CI95%** | **P-FDR** | **Bf_10_** |
| --- | --- | --- | --- | --- | --- | --- | --- | --- | --- |
| Cap vs Cap-MC | A | GEV (proportion) | 0.05 | 0.07 | -4.33 | 0.46 | [-0.02, -0.01] | **< 0.001** | >100*** |
| Cap vs Cap-MC | A | Mean Duration (ms) | 82.10 | 83.49 | -2.14 | 0.19 | [-2.68, -0.10] | 0.06 | 0.97 |
| Cap vs Cap-MC | A | Occurrence (per s) | 1.89 | 2.06 | -4.94 | 0.41 | [-0.24, -0.10] | **< 0.001** | >100*** |
| Cap vs Cap-MC | A | Time Coverage (%) | 15.75 | 17.43 | -4.29 | 0.37 | [-2.46, -0.90] | **< 0.001** | >100*** |
| Cap vs Cap-MC | B | GEV (proportion) | 0.06 | 0.07 | -0.71 | 0.08 | [-0.01, 0.00] | 0.53 | 0.14 |
| Cap vs Cap-MC | B | Mean Duration (ms) | 83.52 | 82.16 | 2.14 | 0.19 | [0.10, 2.61] | 0.06 | 0.96 |
| Cap vs Cap-MC | B | Occurrence (per s) | 1.93 | 1.99 | -2.10 | 0.17 | [-0.12, 0.00] | 0.06 | 0.88 |
| Cap vs Cap-MC | B | Time Coverage (%) | 16.31 | 16.49 | -0.54 | 0.04 | [-0.86, 0.49] | 0.62 | 0.13 |
| Cap vs Cap-MC | C | GEV (proportion) | 0.18 | 0.16 | 3.80 | 0.29 | [0.01, 0.03] | **< 0.001** | 80.01*** |
| Cap vs Cap-MC | C | Mean Duration (ms) | 101.78 | 92.95 | 5.94 | 0.53 | [5.88, 11.77] | **< 0.001** | >100*** |
| Cap vs Cap-MC | C | Occurrence (per s) | 2.54 | 2.40 | 4.93 | 0.36 | [0.08, 0.19] | **< 0.001** | >100*** |
| Cap vs Cap-MC | C | Time Coverage (%) | 26.32 | 22.64 | 6.68 | 0.50 | [2.58, 4.77] | **< 0.001** | >100*** |
| Cap vs Cap-MC | D | GEV (proportion) | 0.10 | 0.12 | -3.66 | 0.36 | [-0.02, -0.01] | **< 0.001** | 50.23*** |
| Cap vs Cap-MC | D | Mean Duration (ms) | 91.19 | 91.06 | 0.20 | 0.02 | [-1.17, 1.43] | 0.84 | 0.11 |
| Cap vs Cap-MC | D | Occurrence (per s) | 2.43 | 2.49 | -2.30 | 0.20 | [-0.12, -0.01] | **0.04** | 1.34* |
| Cap vs Cap-MC | D | Time Coverage (%) | 22.30 | 22.84 | -1.37 | 0.12 | [-1.31, 0.24] | 0.21 | 0.27 |
| Cap vs Cap-MC | E | GEV (proportion) | 0.07 | 0.08 | -3.83 | 0.38 | [-0.02, -0.01] | **< 0.001** | 86.24*** |
| Cap vs Cap-MC | E | Mean Duration (ms) | 88.21 | 89.27 | -1.64 | 0.13 | [-2.34, 0.22] | 0.13 | 0.40 |
| Cap vs Cap-MC | E | Occurrence (per s) | 2.16 | 2.29 | -4.50 | 0.36 | [-0.18, -0.07] | **< 0.001** | >100*** |
| Cap vs Cap-MC | E | Time Coverage (%) | 19.32 | 20.60 | -3.78 | 0.28 | [-1.95, -0.61] | **< 0.001** | 73.58*** |
| nCBP vs nCBP-MC | A | GEV (proportion) | 0.06 | 0.06 | 0.55 | 0.09 | [-0.01, 0.01] | 0.76 | 0.28 |
| nCBP vs nCBP-MC | A | Mean Duration (ms) | 88.29 | 85.16 | 2.98 | 0.51 | [1.05, 5.20] | 0.06 | 0.24 |
| nCBP vs nCBP-MC | A | Occurrence (per s) | 1.93 | 1.91 | 0.26 | 0.04 | [-0.13, 0.16] | 0.84 | 0.20 |
| nCBP vs nCBP-MC | A | Time Coverage (%) | 17.05 | 16.36 | 0.96 | 0.17 | [-0.73, 2.11] | 0.52 | 0.19 |
| nCBP vs nCBP-MC | B | GEV (proportion) | 0.10 | 0.11 | -2.24 | 0.38 | [-0.03, 0.00] | 0.09 | 0.64 |
| nCBP vs nCBP-MC | B | Mean Duration (ms) | 91.18 | 91.14 | 0.02 | 0.00 | [-3.77, 3.85] | 0.98 | 1.21* |
| nCBP vs nCBP-MC | B | Occurrence (per s) | 2.05 | 2.21 | -2.30 | 0.39 | [-0.29, -0.02] | 0.09 | 0.54 |
| nCBP vs nCBP-MC | B | Time Coverage (%) | 18.99 | 20.22 | -1.45 | 0.24 | [-2.91, 0.45] | 0.33 | 0.69 |
| nCBP vs nCBP-MC | C | GEV (proportion) | 0.19 | 0.20 | -0.51 | 0.09 | [-0.04, 0.02] | 0.76 | 0.43 |
| nCBP vs nCBP-MC | C | Mean Duration (ms) | 100.07 | 101.28 | -0.27 | 0.05 | [-9.96, 7.54] | 0.84 | 0.20 |
| nCBP vs nCBP-MC | C | Occurrence (per s) | 2.15 | 2.32 | -3.09 | 0.53 | [-0.29, -0.06] | **0.03** | 13.16*** |
| nCBP vs nCBP-MC | C | Time Coverage (%) | 21.97 | 23.94 | -1.40 | 0.24 | [-4.76, 0.81] | 0.33 | 0.38 |
| nCBP vs nCBP-MC | D | GEV (proportion) | 0.12 | 0.10 | 2.32 | 0.39 | [0.00, 0.03] | 0.09 | 3.53** |
| nCBP vs nCBP-MC | D | Mean Duration (ms) | 95.85 | 91.85 | 2.50 | 0.43 | [0.83, 7.17] | 0.09 | 0.41 |
| nCBP vs nCBP-MC | D | Occurrence (per s) | 2.36 | 2.30 | 0.99 | 0.17 | [-0.07, 0.20] | 0.52 | 0.42 |
| nCBP vs nCBP-MC | D | Time Coverage (%) | 22.82 | 21.28 | 1.74 | 0.30 | [-0.22, 3.30] | 0.24 | 0.52 |
| nCBP vs nCBP-MC | E | GEV (proportion) | 0.07 | 0.08 | -0.61 | 0.10 | [-0.01, 0.01] | 0.76 | 1.47* |
| nCBP vs nCBP-MC | E | Mean Duration (ms) | 90.71 | 88.10 | 1.59 | 0.27 | [-0.64, 5.86] | 0.29 | 0.85 |
| nCBP vs nCBP-MC | E | Occurrence (per s) | 2.08 | 2.04 | 0.44 | 0.07 | [-0.14, 0.21] | 0.78 | 0.34 |
| nCBP vs nCBP-MC | E | Time Coverage (%) | 19.17 | 18.20 | 0.97 | 0.16 | [-1.01, 2.95] | 0.52 | 0.47 |
| CBP vs CBP-MC | A | GEV (proportion) | 0.08 | 0.09 | -0.91 | 0.16 | [-0.02, 0.01] | 0.45 | 0.21 |
| CBP vs CBP-MC | A | Mean Duration (ms) | 86.33 | 85.35 | 0.66 | 0.12 | [-1.97, 3.93] | 0.60 | 9.63 |
| CBP vs CBP-MC | A | Occurrence (per s) | 1.96 | 1.98 | -0.23 | 0.04 | [-0.15, 0.12] | 0.87 | 0.19 |
| CBP vs CBP-MC | A | Time Coverage (%) | 17.06 | 17.04 | 0.03 | 0.00 | [-1.52, 1.56] | 0.98 | 0.28 |
| CBP vs CBP-MC | B | GEV (proportion) | 0.08 | 0.07 | 1.63 | 0.28 | [0.00, 0.02] | 0.30 | 1.76* |
| CBP vs CBP-MC | B | Mean Duration (ms) | 90.62 | 87.32 | 2.02 | 0.36 | [0.06, 6.54] | 0.23 | 0.18 |
| CBP vs CBP-MC | B | Occurrence (per s) | 2.10 | 1.95 | 1.51 | 0.26 | [-0.05, 0.35] | 0.31 | 1.98* |
| CBP vs CBP-MC | B | Time Coverage (%) | 19.28 | 17.39 | 1.68 | 0.29 | [-0.34, 4.12] | 0.30 | 0.48 |
| CBP vs CBP-MC | C | GEV (proportion) | 0.16 | 0.15 | 1.34 | 0.23 | [-0.01, 0.04] | 0.31 | 0.21 |
| CBP vs CBP-MC | C | Mean Duration (ms) | 97.95 | 98.75 | -0.22 | 0.04 | [-8.01, 6.41] | 0.87 | 0.19 |
| CBP vs CBP-MC | C | Occurrence (per s) | 2.24 | 2.39 | -2.17 | 0.38 | [-0.29, -0.01] | 0.23 | 3.16** |
| CBP vs CBP-MC | C | Time Coverage (%) | 22.35 | 24.06 | -1.23 | 0.21 | [-4.48, 1.05] | 0.32 | 0.45 |
| CBP vs CBP-MC | D | GEV (proportion) | 0.11 | 0.15 | -3.73 | 0.63 | [-0.06, -0.02] | **0.005** | 86.53*** |
| CBP vs CBP-MC | D | Mean Duration (ms) | 93.18 | 97.16 | -1.29 | 0.22 | [-10.09, 2.12] | 0.31 | 3.04* |
| CBP vs CBP-MC | D | Occurrence (per s) | 2.24 | 2.33 | -1.31 | 0.24 | [-0.24, 0.05] | 0.31 | 0.29 |
| CBP vs CBP-MC | D | Time Coverage (%) | 21.22 | 23.10 | -1.48 | 0.26 | [-4.41, 0.63] | 0.31 | 0.72 |
| CBP vs CBP-MC | E | GEV (proportion) | 0.08 | 0.07 | 2.13 | 0.40 | [0.00, 0.03] | 0.23 | 0.22 |
| CBP vs CBP-MC | E | Mean Duration (ms) | 90.78 | 87.71 | 1.82 | 0.34 | [-0.29, 6.43] | 0.29 | 0.58 |
| CBP vs CBP-MC | E | Occurrence (per s) | 2.18 | 2.06 | 1.13 | 0.20 | [-0.09, 0.32] | 0.35 | 0.20 |
| CBP vs CBP-MC | E | Time Coverage (%) | 20.09 | 18.40 | 1.40 | 0.26 | [-0.70, 4.07] | 0.31 | 0.28 |

Cap, capsaicin pain; nCBP, non-chronic back pain; CBP, chronic back pain; MC, matched control; BF, Bayes factors. Marked bold indicating FDR-corrected significance levels as < 0.05 *** BF_10_ > 10, ** BF_10_ > 3, and * BF_10_ > 1 indicate anecdotal, moderate, and strong evidence in favour of an effect, respectively.

**Table S5.** Comparison differences of temporal microstate differences across pain groups

| **Microstate** | **Metric** | **Contrast** | **Estimate difference of differences** | **z or t value** | **P-FDR** |
| --- | --- | --- | --- | --- | --- |
| A | GEV (proportion) | CBP minus Cap difference | 0.01 | 1.47 | 0.22 |
| A | GEV (proportion) | nCBP minus Cap difference | 0.02 | 3.45 | **0.004** |
| A | GEV (proportion) | nCBP minus CBP difference | 0.01 | 1.07 | 0.35 |
| A | Mean Duration (ms) | CBP minus Cap difference | 2.37 | 1.47 | 0.22 |
| A | Mean Duration (ms) | nCBP minus Cap difference | 4.52 | 3.68 | **0.002** |
| A | Mean Duration (ms) | nCBP minus CBP difference | 2.15 | 1.19 | 0.32 |
| A | Occurrence (per s) | CBP minus Cap difference | 0.15 | 2.05 | 0.08 |
| A | Occurrence (per s) | nCBP minus Cap difference | 0.19 | 2.36 | 0.07 |
| A | Occurrence (per s) | nCBP minus CBP difference | 0.03 | 0.34 | 0.78 |
| A | Time Coverage (%) | CBP minus Cap difference | 1.70 | 1.97 | 0.10 |
| A | Time Coverage (%) | nCBP minus Cap difference | 2.37 | 2.92 | **0.02** |
| A | Time Coverage (%) | nCBP minus CBP difference | 0.67 | 0.64 | 0.61 |
| B | GEV (proportion) | CBP minus Cap difference | 0.01 | 1.78 | 0.13 |
| B | GEV (proportion) | nCBP minus Cap difference | -0.01 | -1.67 | 0.16 |
| B | GEV (proportion) | nCBP minus CBP difference | -0.02 | -2.77 | 0.06 |
| B | Mean Duration (ms) | CBP minus Cap difference | 1.95 | 1.12 | 0.33 |
| B | Mean Duration (ms) | nCBP minus Cap difference | -1.31 | -0.65 | 0.61 |
| B | Mean Duration (ms) | nCBP minus CBP difference | -3.26 | -1.30 | 0.28 |
| B | Occurrence (per s) | CBP minus Cap difference | 0.21 | 2.07 | 0.08 |
| B | Occurrence (per s) | nCBP minus Cap difference | -0.09 | -1.24 | 0.30 |
| B | Occurrence (per s) | nCBP minus CBP difference | -0.30 | -2.56 | 0.06 |
| B | Time Coverage (%) | CBP minus Cap difference | 2.07 | 1.78 | 0.13 |
| B | Time Coverage (%) | nCBP minus Cap difference | -1.05 | -1.15 | 0.32 |
| B | Time Coverage (%) | nCBP minus CBP difference | -3.12 | -2.23 | 0.07 |
| C | GEV (proportion) | CBP minus Cap difference | -0.01 | -0.37 | 0.77 |
| C | GEV (proportion) | nCBP minus Cap difference | -0.03 | -1.77 | 0.13 |
| C | GEV (proportion) | nCBP minus CBP difference | -0.03 | -1.23 | 0.30 |
| C | Mean Duration (ms) | CBP minus Cap difference | -9.62 | -2.47 | 0.06 |
| C | Mean Duration (ms) | nCBP minus Cap difference | -10.03 | -2.16 | 0.08 |
| C | Mean Duration (ms) | nCBP minus CBP difference | -0.41 | -0.07 | 0.94 |
| C | Occurrence (per s) | CBP minus Cap difference | -0.28 | -3.85 | **0.001** |
| C | Occurrence (per s) | nCBP minus Cap difference | -0.31 | -4.94 | **<0.001** |
| C | Occurrence (per s) | nCBP minus CBP difference | -0.03 | -0.31 | 0.79 |
| C | Time Coverage (%) | CBP minus Cap difference | -5.39 | -3.62 | **0.002** |
| C | Time Coverage (%) | nCBP minus Cap difference | -5.65 | -3.76 | **0.002** |
| C | Time Coverage (%) | nCBP minus CBP difference | -0.26 | -0.13 | 0.91 |
| D | GEV (proportion) | CBP minus Cap difference | -0.03 | -2.36 | 0.07 |
| D | GEV (proportion) | nCBP minus Cap difference | 0.03 | 3.82 | **0.001** |
| D | GEV (proportion) | nCBP minus CBP difference | 0.05 | 4.43 | **<0.001** |
| D | Mean Duration (ms) | CBP minus Cap difference | -4.11 | -1.32 | 0.28 |
| D | Mean Duration (ms) | nCBP minus Cap difference | 3.87 | 2.25 | 0.07 |
| D | Mean Duration (ms) | nCBP minus CBP difference | 7.99 | 2.32 | 0.07 |
| D | Occurrence (per s) | CBP minus Cap difference | -0.03 | -0.38 | 0.77 |
| D | Occurrence (per s) | nCBP minus Cap difference | 0.13 | 1.82 | 0.13 |
| D | Occurrence (per s) | nCBP minus CBP difference | 0.16 | 1.65 | 0.16 |
| D | Time Coverage (%) | CBP minus Cap difference | -1.35 | -1.03 | 0.37 |
| D | Time Coverage (%) | nCBP minus Cap difference | 2.08 | 2.15 | 0.08 |
| D | Time Coverage (%) | nCBP minus CBP difference | 3.43 | 2.23 | 0.07 |
| E | GEV (proportion) | CBP minus Cap difference | 0.03 | 3.43 | **0.004** |
| E | GEV (proportion) | nCBP minus Cap difference | 0.01 | 1.16 | 0.32 |
| E | GEV (proportion) | nCBP minus CBP difference | -0.02 | -2.06 | 0.08 |
| E | Mean Duration (ms) | CBP minus Cap difference | 4.13 | 2.30 | 0.07 |
| E | Mean Duration (ms) | nCBP minus Cap difference | 3.67 | 2.09 | 0.08 |
| E | Mean Duration (ms) | nCBP minus CBP difference | -0.46 | -0.20 | 0.87 |
| E | Occurrence (per s) | CBP minus Cap difference | 0.24 | 2.28 | 0.07 |
| E | Occurrence (per s) | nCBP minus Cap difference | 0.16 | 1.80 | 0.13 |
| E | Occurrence (per s) | nCBP minus CBP difference | -0.08 | -0.58 | 0.64 |
| E | Time Coverage (%) | CBP minus Cap difference | 2.97 | 2.40 | 0.07 |
| E | Time Coverage (%) | nCBP minus Cap difference | 2.25 | 2.14 | 0.08 |
| E | Time Coverage (%) | nCBP minus CBP difference | -0.72 | -0.46 | 0.73 |

Cap, capsaicin pain; nCBP, non-chronic back pain; CBP, chronic back pain; MC, matched control. Marked bold indicating FDR-corrected significance levels as < 0.05

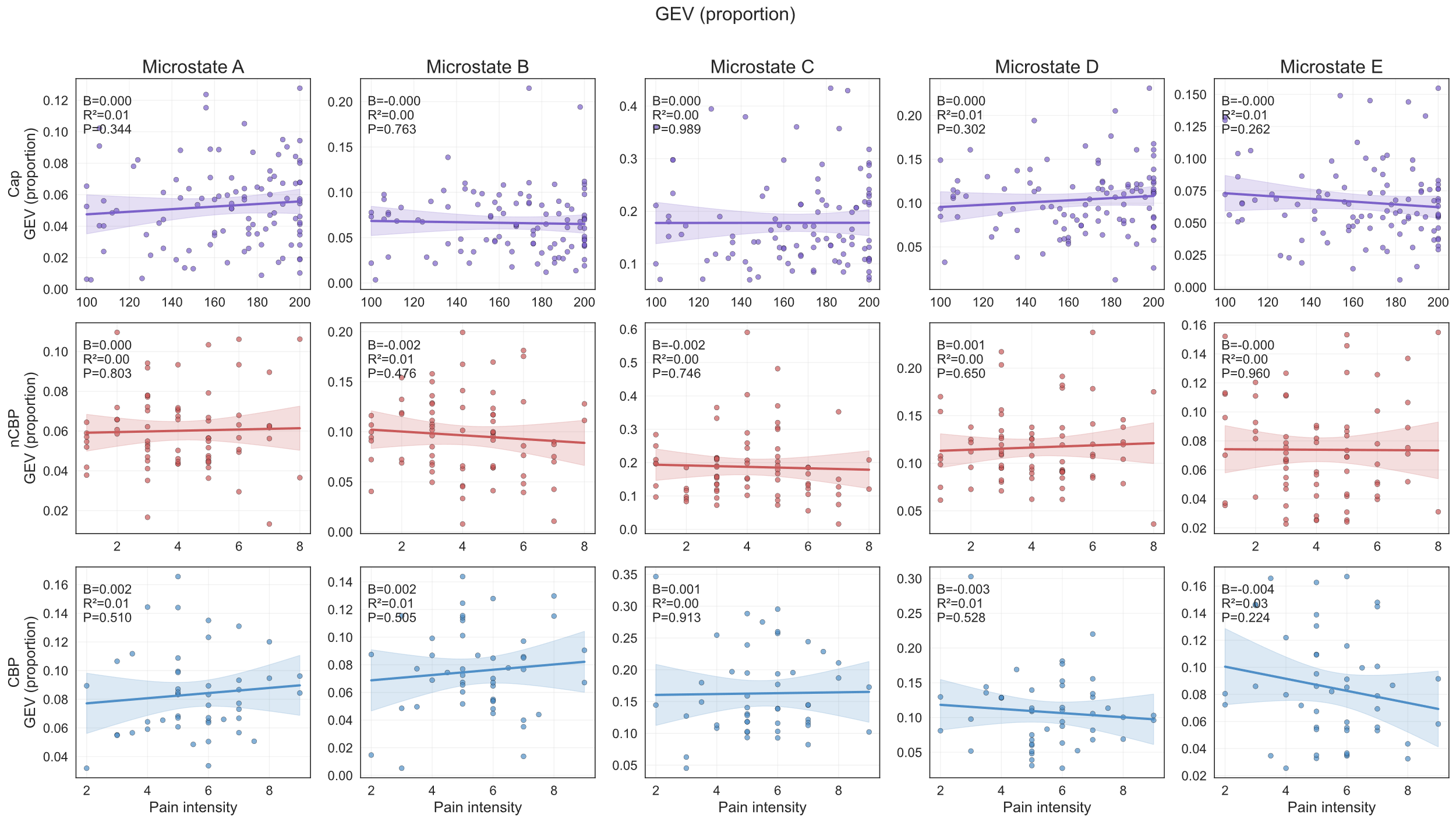

**Figure S2. Associations between pain intensity and global explained variance (GEV) across pain cohorts.** Scatterplots show the relationships between self-reported pain intensity and temporal parameters of microstates A–E within the capsaicin pain (Cap; purple), non-chronic back pain (nCBP; red), and chronic back pain (CBP; blue) cohorts. Solid lines indicate the fitted ordinary least-squares regression relationship, and shaded regions represent the corresponding 95% confidence intervals for the estimated mean response. Pain intensity was expressed on a 0–10 scale for non-chronic and chronic back pain and on a 0-200 for capsaicin pain. No significant associations were observed, after correction across the 20 metric-by-microstate models within each cohort. B is the unstandardized regression coefficient, or slope. R^2^ is the proportion of variance in the microstate metric explained by pain intensity. P is significant value after FDR correction.

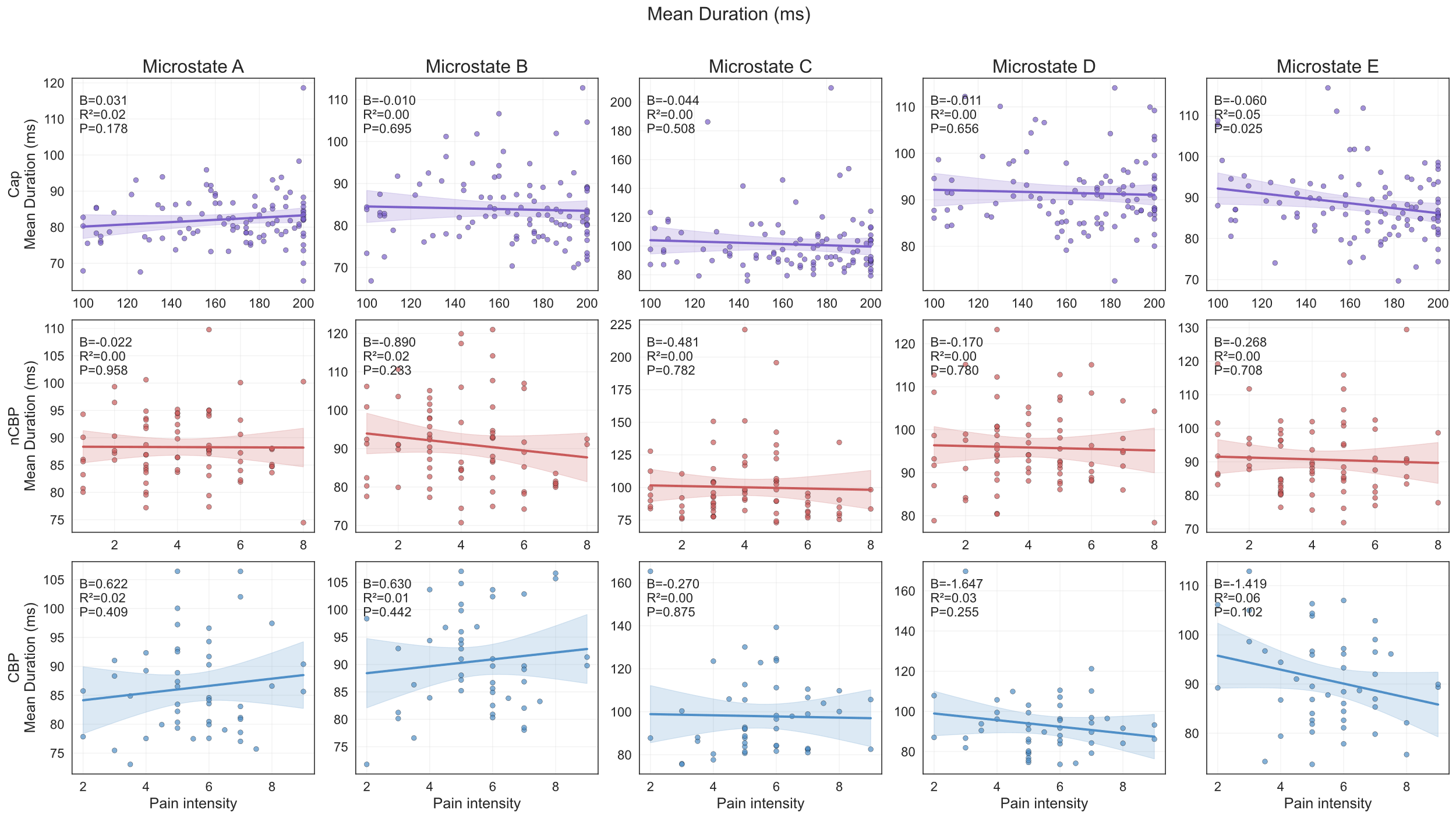

**Figure S3. Associations between pain intensity and mean duration across pain cohorts.** Scatterplots show the relationships between self-reported pain intensity and temporal parameters of microstates A–E within the capsaicin pain (Cap; purple), non-chronic back pain (nCBP; red), and chronic back pain (CBP; blue) cohorts. Each point represents one participant. Solid lines indicate the fitted ordinary least-squares regression relationship, and shaded regions represent the corresponding 95% confidence intervals for the estimated mean response. Pain intensity was expressed on a 0–10 scale for non-chronic and chronic back pain and on a 0-200 for capsaicin pain. Significant association was observed between pain intensity and microstate E in Cap group. B is the unstandardized regression coefficient, or slope. R^2^ is the proportion of variance in the microstate metric explained by pain intensity. P is significant value after FDR correction.

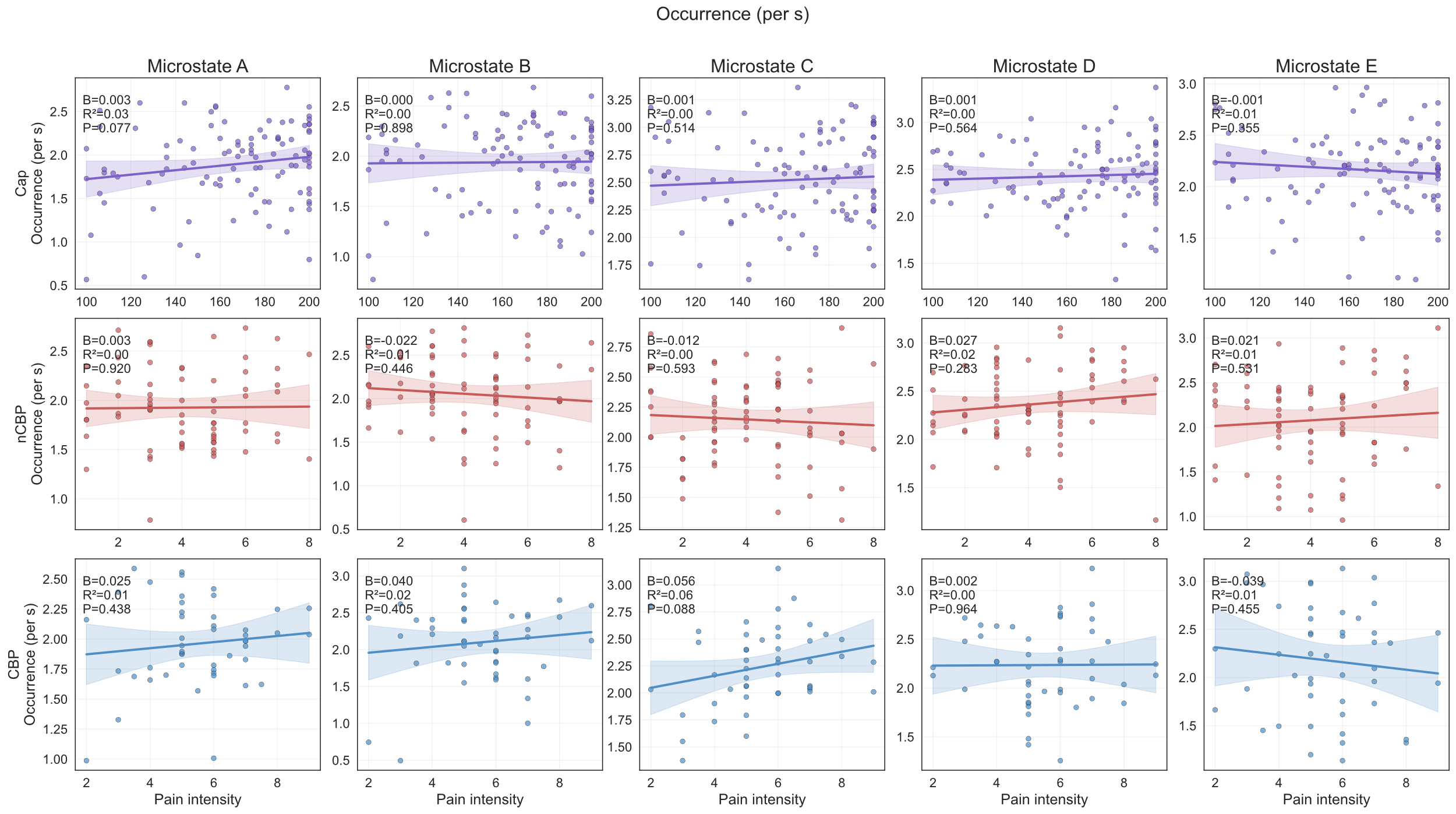

**Figure S4. Associations between pain intensity and occurrence across pain cohorts.** Scatterplots show the relationships between self-reported pain intensity and temporal parameters of microstates A–E within the capsaicin pain (Cap; purple), non-chronic back pain (nCBP; red), and chronic back pain (CBP; blue) cohorts. Each point represents one participant. Solid lines indicate the fitted ordinary least-squares regression relationship, and shaded regions represent the corresponding 95% confidence intervals for the estimated mean response. Pain intensity was expressed on a 0–10 scale for non-chronic and chronic back pain and on a 0-200 for capsaicin pain. No significant associations were observed, after correction across the 20 metric-by-microstate models within each cohort. B is the unstandardized regression coefficient, or slope. R^2^ is the proportion of variance in the microstate metric explained by pain intensity. P is significant value after FDR correction.

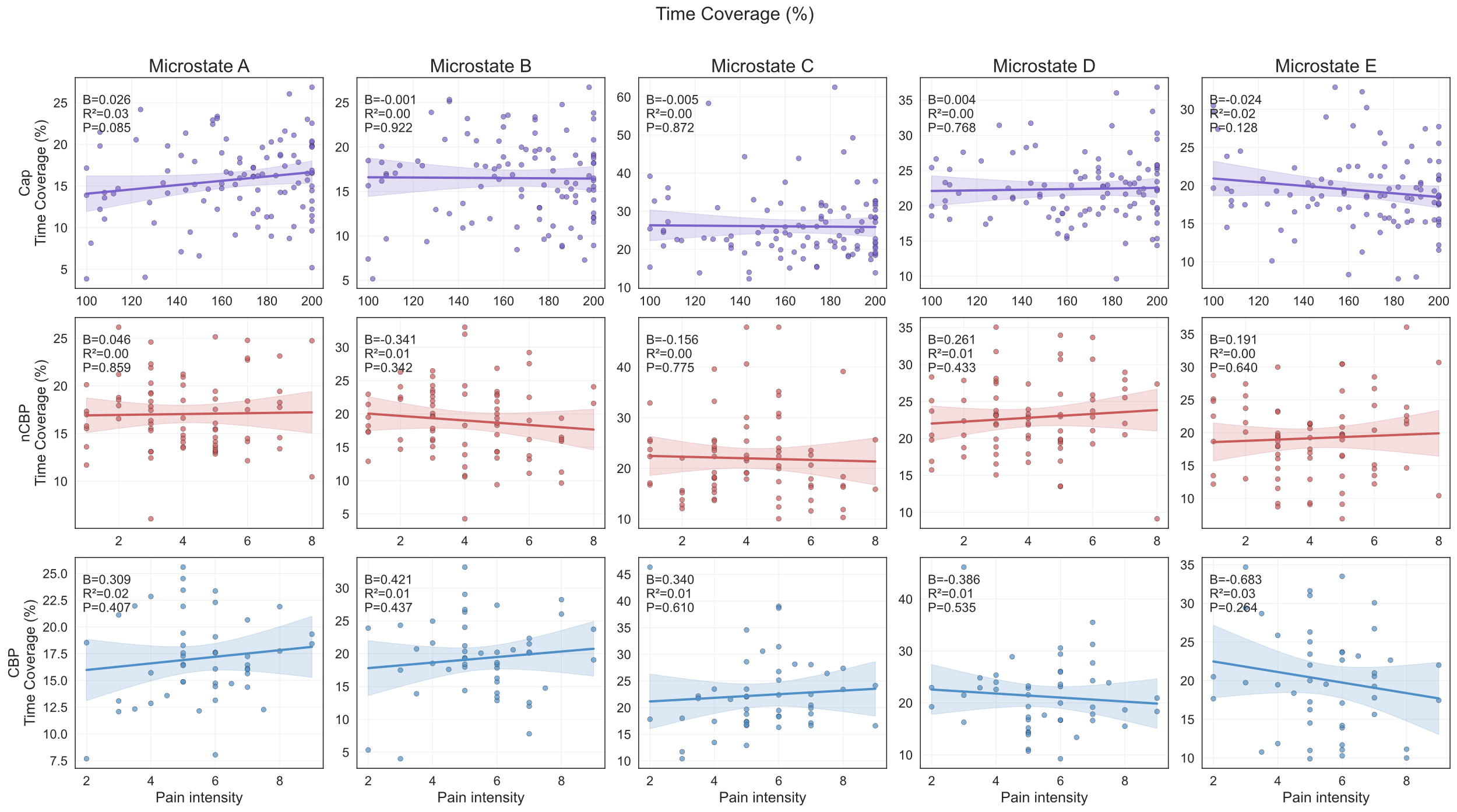

**Figure S5. Associations between pain intensity and time coverage across pain cohorts.** Scatterplots show the relationships between self-reported pain intensity and temporal parameters of microstates A–E within the capsaicin pain (Cap; purple), non-chronic back pain (nCBP; red), and chronic back pain (CBP; blue) cohorts. Each point represents one participant. Solid lines indicate the fitted ordinary least-squares regression relationship, and shaded regions represent the corresponding 95% confidence intervals for the estimated mean response. Pain intensity was expressed on a 0–10 scale for non-chronic and chronic back pain and on a 0-200 for capsaicin pain. No significant associations were observed, after correction across the 20 metric-by-microstate models within each cohort. B is the unstandardized regression coefficient, or slope. R^2^ is the proportion of variance in the microstate metric explained by pain intensity. P is significant value after FDR correction.

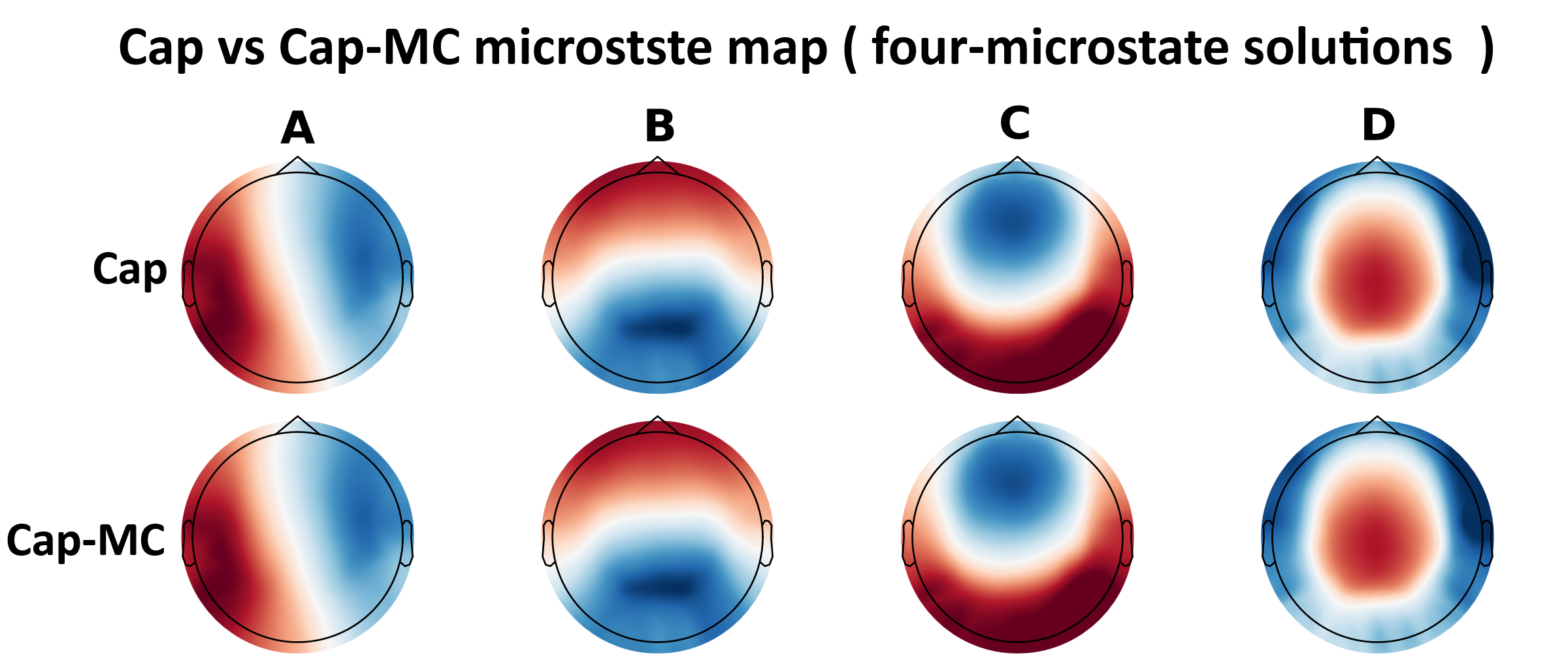

**Figure S6. Group-level topographies of the four-microstate solutions for the sensitivity comparison of capsaicin pain and matched condition.** Microstate maps A-D are shown for capsaicin pain group and its matched condition. Microstate labels were assigned based on their correspondence with commonly reported canonical microstate clusters. Cap, capsaicin pain; MC, matched condition

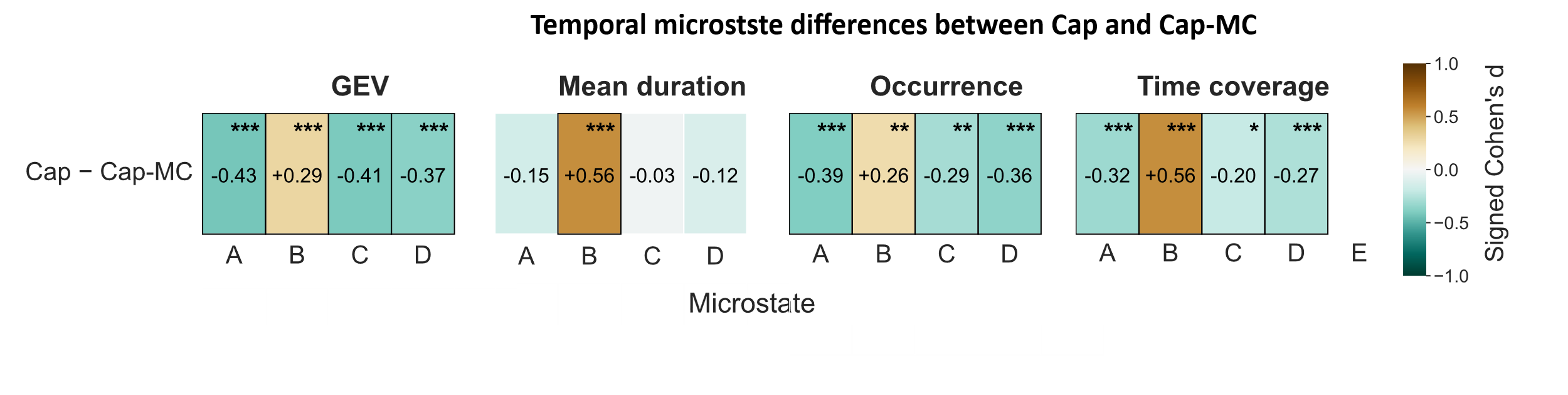

**Figure S7. Temporal microstate differences of the sensitivity comparison between capsaicin pain and matched condition. A:** Standardized differences in global explained variance, mean duration, occurrence, and time coverage between capsaicin pain group and its matched condition. Values represent signed Cohen’s d, with positive values indicating higher metric values in the pain group and negative values indicating lower values relative to the matched condition. Paired-samples tests were used for the capsaicin comparison. Cap, capsaicin pain; MC, matched condition

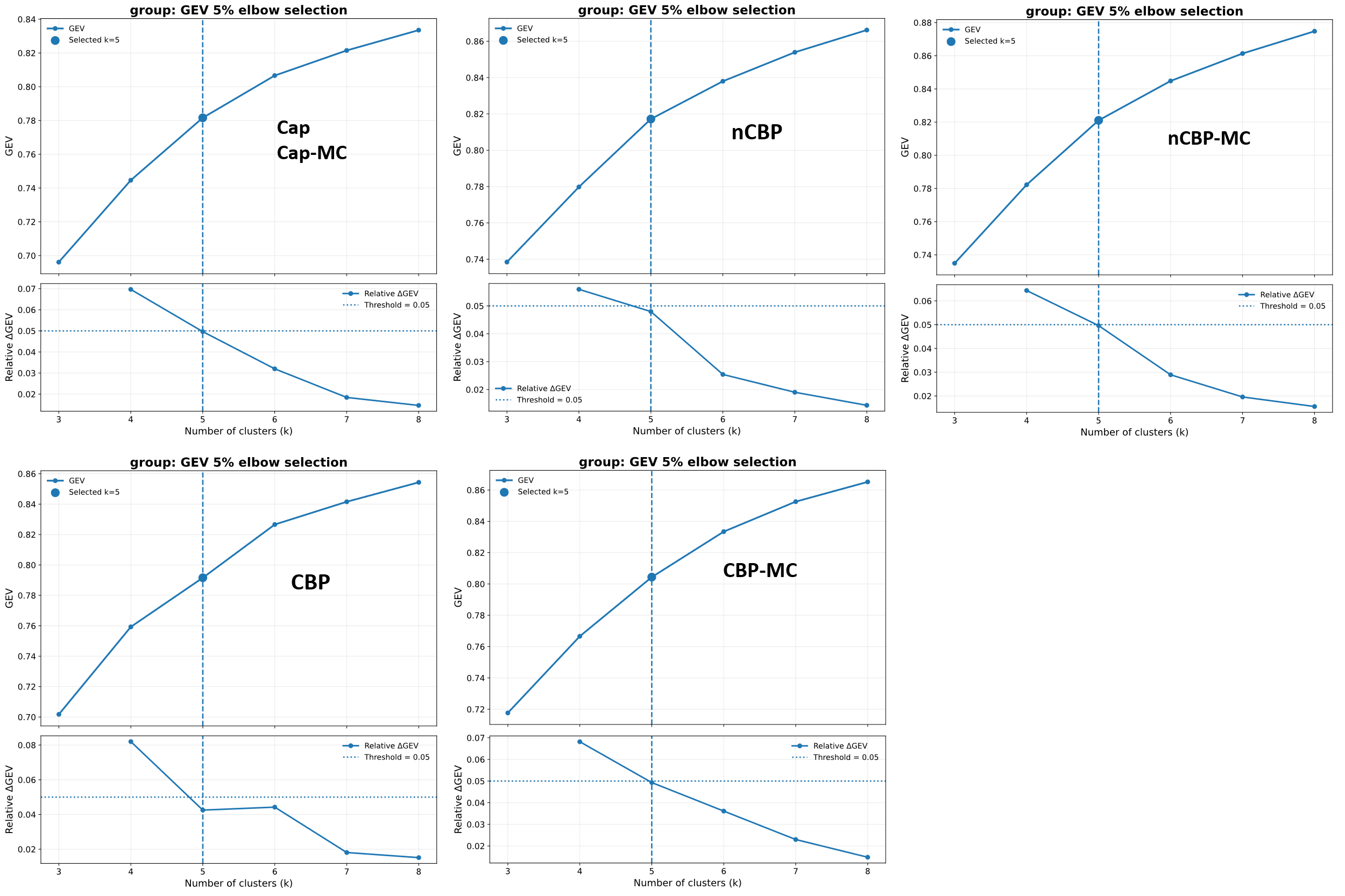

**Figure S8. Global explained variance elbow for detecting the number of microstates.** Five-microstate solution was detected by GEV elbow (the increase in GEV for one number of microstates relative to the previous number of microstates was <5%) as the most appropriate cluster. Cap, capsaicin pain; nCBP, non-chronic back pain; CBP, chronic back pain; MC, matched control. GEV, Global explained variance

**Table S6.** Group-level global explained variance indices

| **Group** | **Cluster** | | | | | |
| --- | --- | --- | --- | --- | --- | --- |
|  | **3** | **4** | **5** | **6** | **7** | **8** |
| Cap | 0.70 | 0.74 | 0.78 | 0.81 | 0.82 | 0.83 |
| Cap-MC | 0.70 | 0.74 | 0.78 | 0.81 | 0.82 | 0.83 |
| nCBP | 0.74 | 0.78 | 0.82 | 0.84 | 0.85 | 0.87 |
| nCBP-MC | 0.73 | 0.78 | 0.82 | 0.84 | 0.86 | 0.87 |
| CBP | 0.70 | 0.76 | 0.79 | 0.83 | 0.84 | 0.85 |
| CBP-MC | 0.72 | 0.77 | 0.80 | 0.83 | 0.85 | 0.87 |

Cap, capsaicin pain; nCBP, non-chronic back pain; CBP, chronic back pain; MC, matched control
